# Cell cycle and temporal transcription factors regulate proliferation and neuronal diversity of dedifferentiation-derived neural stem cells

**DOI:** 10.1101/2022.07.24.501087

**Authors:** Kellie Veen, Francesca Froldi, Qian Dong, Edel Alvarez-Ochoa, Phuong-Khanh Nguyen, Kieran F Harvey, John P D McMullen, Owen Marshall, Patricia R Jusuf, Louise Y Cheng

**Affiliations:** Peter MacCallum Cancer Centre, Melbourne, VIC 3000, Australia; Sir Peter MacCallum Department of Oncology, The University of Melbourne, VIC 3010, Australia; Department of Anatomy and Physiology, The University of Melbourne, VIC 3010, Australia; School of Biosciences, The University of Melbourne, VIC 3010, Australia; Menzies Institute for Medical Research, Medical Science Precinct, 17 Liverpool Street, Hobart TAS 7000, Australia

## Abstract

Dedifferentiation is the reversion of differentiated cells to a stem cell like fate, whereby, the gene expression program of mature cells is altered and genes associated with multipotency are expressed. Appropriate terminal differentiation of NSCs is essential for restricting the overall number of neurons produced; in addition, faithful production of neuronal subtypes that populate the brain is important for NSC function. Both characteristics of NSCs are specified through temporal patterning of the NSCs driven by the successive expression of temporal transcription factors (tTFs). In this study, we found that ectopic NSCs induced via bHLH transcription factor Deadpan (Dpn) expression fail to undergo timely expression of temporal transcription factors (tTFs), where they express mid-tTF, Sloppy-paired 1 (Slp-1) and fail to express late-tTF Tailless (Tll); consequently generating an excess of Twin of eyeless (Toy) positive neurons at the expense of Reversed polarity (Repo) positive glial cells. In addition to disrupted production of neuronal/glial progeny, Dpn overexpression also resulted in stalled progression through the cell cycle, and a failure to undergo timely terminal differentiation. Mechanistically, DamID studies demonstrated that Dpn directly binds to both Dichaete (D), a Sox-box transcription factor known to repress Slp-1, as well as a number of cell cycle genes. Promoting cell cycle progression or overexpression of D were able to re-trigger the progression of the temporal series in dedifferentiated NBs, restoring both neuronal diversity and timely NB terminal differentiation.

## Introduction

The central nervous system (CNS) is the cognitive control centre of the body and is generated by neural stem cells (NSCs) during development. *Drosophila* NSCs, called neuroblasts (NBs), exhibit key properties of mammalian NSCs such as self-renewal, temporal control of proliferation, and production of different neural subtypes that will form the adult brain. Thus, NBs serve as a powerful model to study the mechanisms that underlie the determination of overall brain size and neuronal diversity (Harding and White, 2018).

Molecular differences exist between different NB populations located in different parts of the brain. NBs within the ventral nerve cord (VNC) and the central brain (CB) of the larval CNS undergo Type I divisions, whereby the NB divides asymmetrically to produce a NB and a smaller ganglion mother cell (GMC). The GMC then divides to produce two daughter cells, which terminally differentiate into neurons or glia. A subset of NBs in the larval CB undergo Type II division, in which the NBs asymmetric division produces a NB and an intermediate neural progenitor (INP). INPs mature prior to dividing asymmetrically to produce another progenitor and a GMC. Outer proliferation centre (OPC) NBs within the optic lobes (OL) are produced from a pseudostratified neuroepithelium through a differentiation wave characterised by proneural factor expression, termed the ‘proneural wave’ (Egger et al., 2007; Yasugi et al., 2008). These NBs undergo asymmetric division, and are broadly thought to behave similarly to Type I NBs.

In addition to positional identity, NBs can also be temporally defined by the expression pattern of sequential markers, known as the temporal series or temporal transcription factors (tTFs). The tTFs confer temporal identity to neurons born within each tTF window, and their cross-regulation ensures unidirectional progression through the temporal series (Li et al., 2013). Type I, II and medulla NBs all express their unique set of tTFs, however, in all the regions of the CNS, tTFs play conserved roles to control neuronal diversity and proliferative potential of NBs (Abdusselamoglu et al., 2019; Bayraktar and Doe, 2013; Eroglu et al., 2014; Konstantinides et al., 2022; Li et al., 2013; Liu et al., 2015; Maurange et al., 2008; Pahl et al., 2019; Ren et al., 2017; Suzuki et al., 2013; Syed et al., 2017; Zhu et al., 2022).

One of the key regulatory mechanisms of brain size and neuronal diversity is through control of NB identity via cell fate maintenance. We and others have shown that a number of transcription factors including Longitudinals Lacking (Lola), Midlife crisis and Nervous fingers 1 (Nerfin-1) prevent neuron-to-NB reversion by repressing the activity of NB and cell-cycle genes in post-mitotic neurons in the OLs (Carney et al., 2013; Froldi et al., 2015; Froldi et al., 2015; Froldi et al., 2019; Southall et al., 2014; Vissers et al., 2018; Xu et al., 2017). Additionally, signalling pathways such as Notch, have been shown to play a role in dedifferentiation, whereby Notch hyperactivation induces ectopic NBs (Vissers et al., 2018; Xu et al., 2017). However, beyond the fact that misexpression of these factors and pathways caused the formation of ectopic NBs, whether these dedifferentiated NBs faithfully produce the correct number and types of neurons or glial cells, or undergo timely terminal differentiation, has not been assessed. These characteristics are key determinants of overall CNS size and function, thus are important parameters when considering whether dedifferentiation leads to tumourigenesis or can be appropriately utilised for regenerative purposes.

In this study, we identified the basic helix-loop-helix (bHLH) transcription factor, Deadpan (Dpn), as a mediator of dedifferentiation downstream of Nerfin-1. Overexpression of Dpn caused the formation of supernumery NBs in deep neuronal layers of the medulla within the OLs. The dedifferentiated NBs immediately acquired and maintained Eyeless (Ey)/Sloppy paired 1 (Slp) positive mid-temporal identity throughout larval neurogenesis, delaying the onset of the late tTF Tailess (Tll). As a result, excessive Toy^+^ progeny was made at the expense of Tll^+^ neurons and Repo^+^ glial cells. Furthermore, ectopic NBs generated via Dpn overexpression were stalled in the cell cycle and failed to undergo terminal differentiation in a timely fashion. Mechanistically, our DamID analysis showed that Slp-1 and 2, group B Sox-transcription factor, Dichaete (D), and cell cycle genes, *string* (*stg*), *cyclin D* (*Cyc D*) and *cyclin E* (*Cyc E*), are direct targets of Dpn. Overexpression of D was able to promote the progression of the temporal series in ectopic NBs generated via Dpn overexpression, rescuing both neuronal diversity and terminal differentiation. Similarly, promoting NB progression through G1/S phase of the cell cycle was able to restore temporal progression, neuronal diversity, and terminal differentiation. Dedifferentiation induced via Notch hyperactivation also exhibited stalled temporal progression, which was restored via D overexpression.

Together, our findings suggest that cell cycle regulators and temporal transcription factors are key determinants of proliferation and termination profiles of dedifferentiated NBs. In order for us to utilise dedifferentiated NSCs for regenerative purposes and to prevent tumorigenesis that often originate from differentiated cell types, we will have to recreate the right temporal profiles and ensure cell cycle progression occurs appropriately.

## Results

### Overexpression of Deadpan, results in ectopic NB formation in the medulla regions of the optic lobes

We and others have previously shown that Nerfin-1 is a transcription factor that maintains neuronal differentiation in Type I, II and medulla neuronal lineages (Froldi et al., 2015; Vissers et al., 2018; Xu et al., 2017). In the absence of Nerfin-1, neurons undergo dedifferentiation to acquire a NB like fate, by turning on bona-fide NB markers such as Deadpan (Dpn) and Miranda (Mira) (Froldi et al., 2015; Vissers et al., 2018; Xu et al., 2017). In the OLs, symmetrically dividing neuroepithelial cells differentiate into medulla NBs located at the superficial CNS surface (0 µm, Figure 1A, 1B, pink), which in turn generates medulla neurons that occupy the deep layers of the CNS at around 8-12 µm from the superficial surface (Figure 1A, Figure 1B, green) (Egger et al., 2007; Yasugi et al., 2008). Pan-neural bHLH transcription factor Dpn was identified to be a Nerfin-1 target gene in our previous DamID analyses (Vissers et al., 2018). To assess whether Dpn functionally lies downstream of Nerfin-1, we tested whether *UAS-dpn* caused dedifferentiation. Using three different neuronal/NB drivers – *GMR31HI08-GAL4* (Jenett et al., 2012), *ey^OK107^-GAL4* (Morante et al., 2011) and *eyR16F10-GAL4* (Jenett et al., 2012), we found that Dpn overexpression in the medulla region of the brain induced the formation of ectopic NBs in deep sections of the CNS, phenocopying the loss of Nerfin-1 (Figure S1A-F, Vissers et al., 2018).

**Figure 1.**
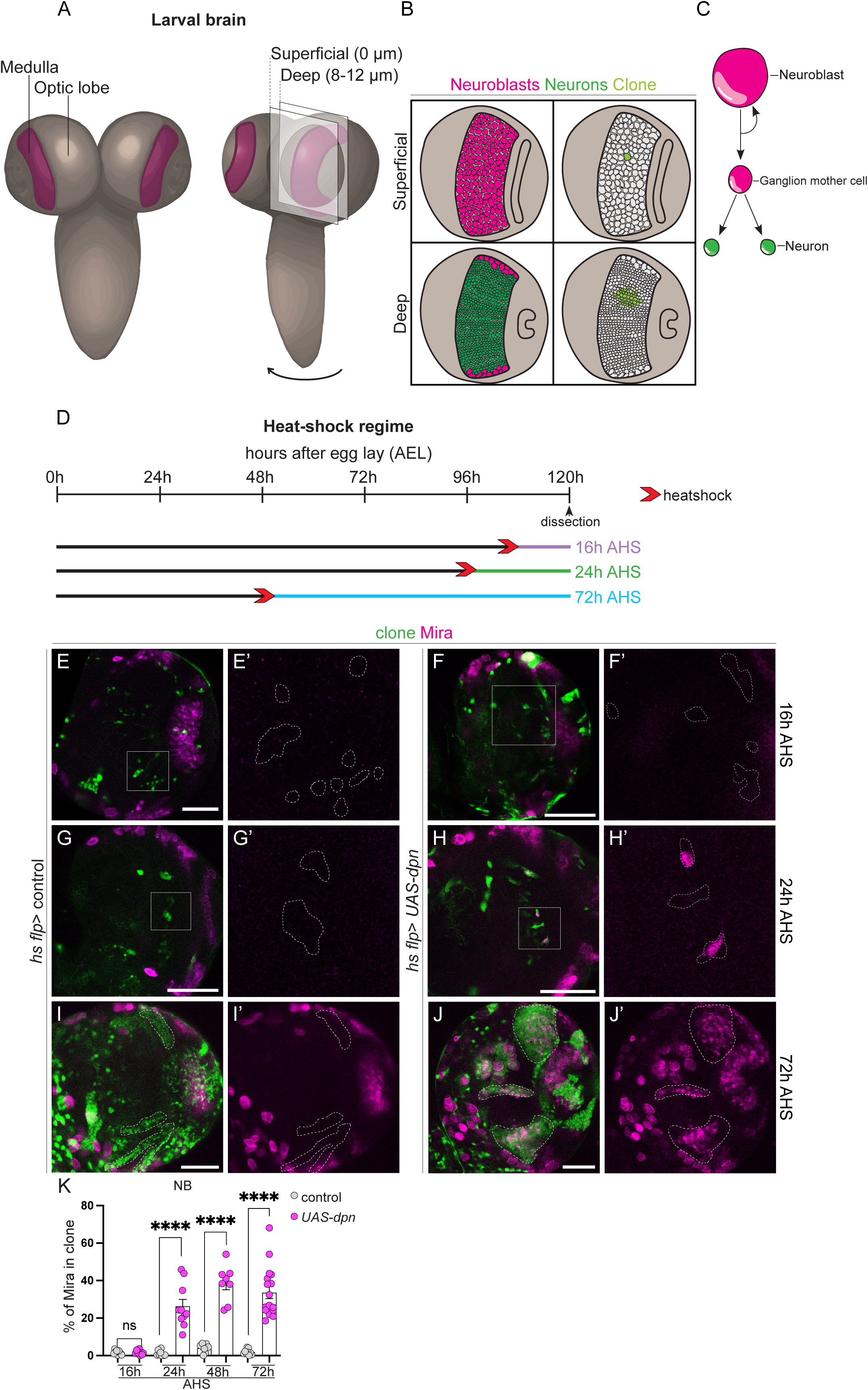
Dpn overexpression causes reversion of neurons to NBs in the deep layers of the developing CNS. (A) Schematic representation of the larval central nervous system (CNS). Medulla (depicted in maroon) is located with the Optic Lobes (OLs) of the CNS. Cross section and cell composition at superficial surface (0 μm) and deep layer (8-12 μm) of the OLs are depicted in B. (B) (left) Superficial surface is occupied by neuroblasts (NBs, pink), and deep layers are occupied by neurons (green). (right) Cross section through a *hs flp* clone (light green) showing the clone consists of a single NB in the superficial layer and its neuronal progeny in the deep layer. (C) Schematic representation of NB divisions: NBs undergo asymmetric division to give rise to a Ganglion mother cell (GMC) which produces two neurons. (D) Schematic depicting the heat shock regimes used in (E-K). Clones were induced via heat shock (red arrows), and dissected at 16 hr (purple), 24 hr (green), and 72 hr (blue) after heat shock. (E’, F’, G’, H’) are the magnified images of the square region outlined in (E, F, G, H). (E-J) are merged images, (E’-J’) are single channel images of (E-J) depicting the presence of NBs (Mira+). Clones (green) are outlined by dotted lines. All images in this figure, and all following figures are single confocal sections. (E-E’ and F-F’) No Mira+ NBs are recovered in deep sections of control or *UAS-dpn* clones at 16 hr after clone induction. (G-G’ and H-H’) At 24 hr after clone induction, Mira+ NBs are recovered in *UAS-dpn* but not control clones. (I-I’and J-J’) At 72h after clone induction, numerous Mira+ NBs are recovered in *UAS-dpn* but not in control clones. (K) Quantification of % Mira+ cells in control and *UAS-dpn* clones (calculated as the ratio of Mira+ cell volume as a percentage of total clone volume). 16 hr (Control, n=9, m=1.584 ±0.3779, *UAS-dpn*, n=9, m=1.657 ±0.37), 24 hr (Control, n=9, m=1.150 ±0.4781, *UAS-dpn*, n=10, m=26.37 ±3.629), 48 hr (Control, n=7, m=3.971 ±0.9249, *UAS-dpn*, n=8, m=38.56 ±3.446), 72 hr (Control, n=9, m=1.874 ±0.4897, *UAS-dpn*, n=17, m=33.62 ±3.243). Data are represented as mean ± SEM. P-values were obtained using unpaired t-test, and Welch’s correction was applied in case of unequal variances. ****p < 0.0001. Scale bars: 50 μm.

Dedifferentiation was also confirmed using clonal analysis(Figure 1D, via heat shock induced actin-Gal4 flp-out). Ectopic NBs (Mira^+^) in clones were first recovered at 24 hrs after heat shock AHS in deep sections of the CNS (Figure 1 E-H’, quantified in Figure 1K), indicating that NBs first generated mature neurons (as indicated by the lack of ectopic NBs at 16 hrs AHS), which then underwent dedifferentiation to give rise to ectopic NBs by 24 hrs AHS. These ectopic NBs did not express neuronal marker Elav (Figure S1H-H’’), and were actively dividing, as indicated by M-phase marker phosphor-Histone H3 (pH3, Figure S1G-G’’). By 72 hrs AHS, clones overexpressing Dpn consisted of multiple Mira^+^ cells (Figure 1I-K). To assess whether these ectopic NBs were capable of generating neuronal progeny, we performed an EdU-pulse-chase experiment, where the animals were fed EdU labelled food for 2 hrs and chased for 2 hrs with EdU-free food (Figure S1I). In 70% of the clones assessed, Mira^+^ cells inherited EdU during the pulse and produced Mira^−^/EdU^+^ progeny during the chase (Figure S1J-K), indicating that dedifferentiated NBs produced differentiated progeny.

As Dpn was identified to be a target of Nerfin-1, we next investigated the epistatic relationship between Dpn and Nerfin-1. We induced clones expressing an RNAi against Nerfin-1 and compared the level of dedifferentiation with that of Nerfin-1 RNAi clones that also expressed Dpn RNAi (Figure S2A-D). We found that the Dpn RNAi expression significantly reduced the number of ectopic NBs induced by Nerfin-1 knockdown, suggesting that Dpn lies downstream of Nerfin-1 to mediate neuronal cell fate maintenance.

### Ectopic neuroblasts generated via Dpn overexpression express mid-temporal transcription factors

NBs contribute to the sequential generation of different neuron types achieved through the changing expression of temporal transcription factors (tTFs) (Apitz and Salecker, 2014). In the OLs, medulla NBs sequentially expressed a series of tTFs including early factor Homothorax (Hth) expressed by young NBs, mid tTFs Eyeless (Ey), Sloppy paired (Slp) and Dichaete (D) expressed by middle-aged NBs, and late tTF Tailless (Tll) expressed by old NBs (Li et al., 2013; Suzuki et al., 2013). The tTFs contribute to the progression of the temporal series by activating the next tTF while repressing the previous tTF (Figure 2A). Medulla NBs continuously transit through the temporal cascade as they age; therefore, at any given time, NBs in the superficial layer of the medulla would express Hth, Ey, Slp, D, Tll or a combination of these tTFs (Figure S2E-I’). As expected, at 72hrs after clone induction (Figure 2B), around 20-30% the superficial NBs were positive for either Hth, Ey, Slp, D or Tll (or a combination of these factors, Figure 2H). Surprisingly, the ectopic NBs in UAS-Dpn clones mostly expressed mid-tTFs Ey, Slp, and D, while very few ectopic NBs expressed the early tTF Hth or late tTF Tll (Figure 2C-G’’, H).

**Figure 2.**
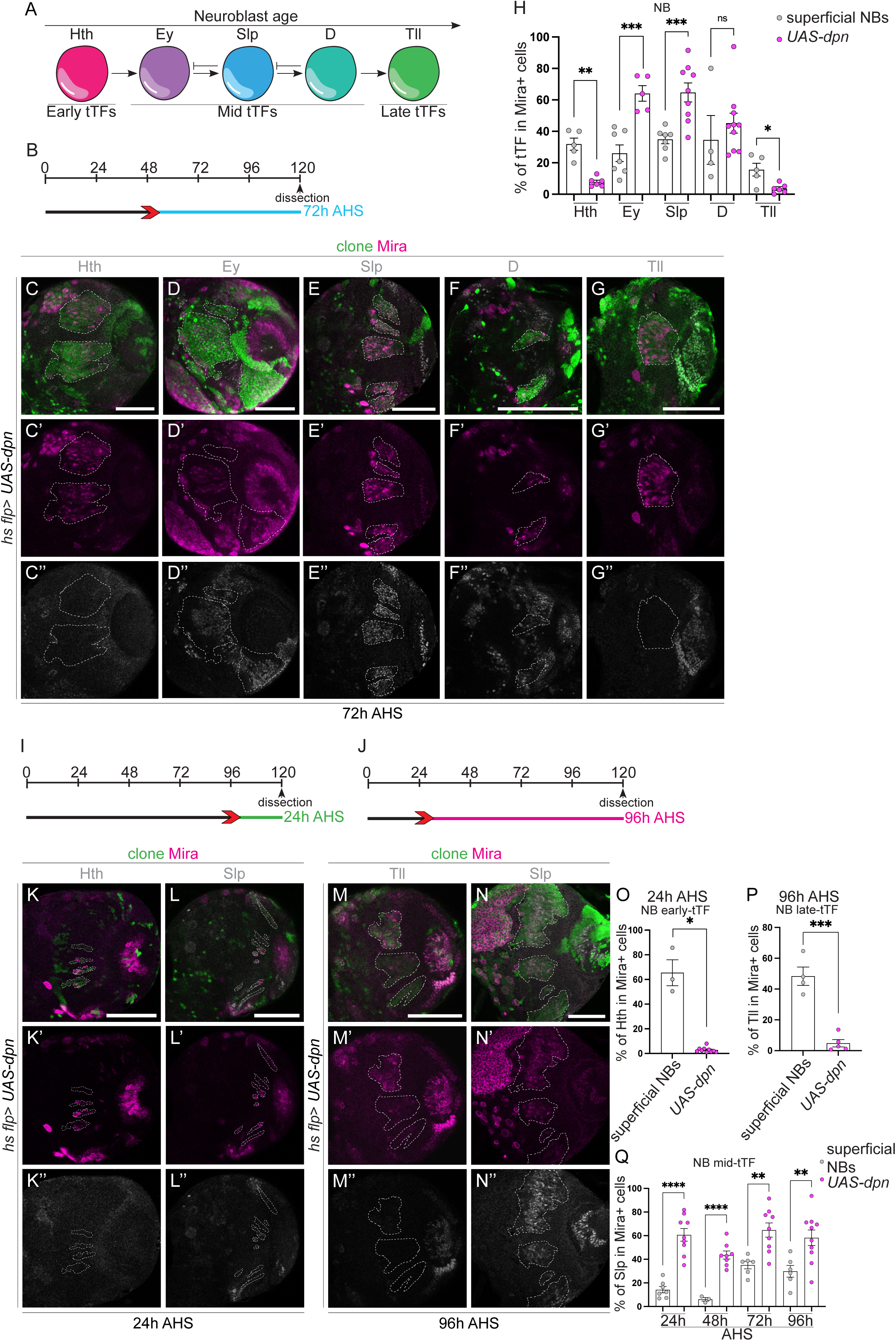
Ectopic NBs generated via *dpn* overexpression express mid-temporal transcription factors. (A) Schematic representation of temporal series in neuroblasts (NBs). NBs express temporal transcription factors (tTFs) (Hth, Ey, Slp, D, Tll) as they age. These can be categorised into early, mid, and late tTFs. (C-G, K-N) Representative images of the deep medulla neuronal layer in the larval optic lobe, in which *UAS-dpn* is driven in clones by *hs flp* (marked by GFP and outlined) and stained with the stem cell marker, Miranda (Mira, magenta), and various temporal transcription factors (tTFs) (grey). (B) Schematic depicting the heat shock regimes used in C-G’’. Clones were induced via heat shock (red arrows) and dissected at 72 hr (blue) after heat shock. (C-G’’) In deep sections of the NB clone induced via *UAS-dpn*, Mira+ NBs do not express early tTF Hth or late tTF Tll, but are positive for mid tTFs Ey, Slp and D. (H) Quantification of volume of cells that express a specific tTF as % of total Mira+ NB volume within a clone. Hth (Control, n= 5, m= 31.86 ±3.89, *UAS-dpn*, n= 6, m= 7.779 ±1.212), Ey (Control, n= 7, m= 26.05 ±5.302, *UAS-dpn*, n= 5, m= 64.08 ±4.936), Slp (Control, n= 7, m= 34.87 ±2.753, *UAS-dpn*, n= 9, m= 64.72 ±6.045), D (Control, n= 4, m=34.57 ±15.49, *UAS-dpn*, n= 10, m= 45.2 ±6.339), Tll (Control, n= 5, m= 15.65 ±4.036, *UAS-dpn*, n= 6, m= 3.584, ±1.87). (I) Schematic depicting the heat shock regimes used in K-L’’. Clones were induced via heat shock (red arrows) and dissected at 24h (green) after heat shock. (K-K’’ and L-L’’) At 24h after clone induction, Mira+ NBs express mid tTF Slp and do not express early tTF Hth, quantified in O and Q. (J) Schematic depicting the heat shock regimes used in M-N’’. Clones were induced via heat shock (red arrows) and dissected at 96h (green) after heat shock. (M-M’’ and N-N’’) At 96h after clone induction, Mira+ NBs express mid tTF Slp and do not express late tTF Tll quantified in P and Q. (O) Quantification of % of Hth+ cells in Mira+ cells in control and *UAS-dpn* clones (expressed as the ratio between Hth+ volume and total Mira+ volume within a clone). Control, n= 3, m= 65.42 ±10.52, *UAS-dpn*, n= 10, m= 2.866 ±0.651 (P) Quantification of % of Tll+ cells in Mira+ cells in control and *UAS-dpn* clones (expressed as the ratio between Tll+ volume and total Mira+ volume within a clone). Control, n= 4, m= 48.34 ±5.956, *UAS-dpn*, n= 5, m= 4.805 ±2.375. (Q) Quantification of % of Slp+ cells in Mira+ cells in control and *UAS-dpn* clones (expressed as the ratio between Slp+ volume and total Mira+ volume within a clone). 24 hr (Control, n= 7, m= 14.25 ±2.696, *UAS-dpn*, n= 9, m= 60.66 ±5.384), 48 hr (Control, n= 3, m= 6.096 ±1.509, *UAS-dpn*, n= 8, m= 43.64 ±3.398), 72 hr (Control, n= 6, m= 35.13 ±3.242, *UAS-dpn*, n= 9, m= 64.72 ±6.045), 96 hr (Control, n= 6, m= 29.85 ±4.951, *UAS-dpn*, n= 10, m= 58.2 ± 6.603). Data are represented as mean ± SEM. P-values were obtained performing unpaired t-test, and Welch’s correction was applied in case of unequal variances. ****p<0.0001, ***p<0.001, **p<0.005, *p<0.05. Scale bars: 50 μm.

As the tumour cell-of-origin can define the competence of tumour NBs to undergo malignancy (Farnsworth et al., 2015; Narbonne-Reveau et al., 2016), we next tested whether the temporal identity of the dedifferentiated NBs were conferred by the age of the neurons they were derived from. At 24 hrs AHS (Figure 2I), the majority of the newly generated control NBs expressed the early tTF Hth (~65%); in contrast, only 2% of Dpn overexpression NBs were Hth^+^ (Figure 2K-L’’, O). Whereas at 96 hrs AHS (Figure 2J), ~50% of the control NBs were positive for the late tTF Tll; in contrast, only 5% of the ectopic NBs generated via Dpn overexpression were Tll^+^ (Figure 2M-N’’, P). At both of these time points (as well as additional time points at 48 hrs and 72 hrs after clone induction), we found that between 40-70% of the ectopic NBs expressed the mid tTF Slp (Figure 2Q). Together, it appears that ectopic NBs immediately adopted a mid-temporal identity shortly after clone induction and continue to do so throughout larval development.

### NBs generated via Dpn overexpression exhibit delayed terminal differentiation

Medulla NBs progress through the series of temporal factors, until the oldest medulla NBs defined by Glial cells missing (Gcm), triggers the transition from neurogenesis to gliogenesis and cell cycle exit (Li et al., 2013; Zhu et al., 2021). The timing of the NB cell cycle exit occurred at around 16 hrs after pupal formation (APF) in control OLs (29°C, Figure 3A-D, J). Upon Dpn overexpression (under the control of *eyR16F10-GAL4* (Jenett et al., 2012)), NBs termination was delayed (Figure 3F-J). *UAS-dpn* NBs maintained the expression of the mid-tTF Slp for much of the pupal neurogenesis (Figure S2 J-K’’’, S3 A-E), but prior to their terminal differentiation, they did begin to reduce Slp and express the late tTF Tll (Figure S3 D-E). Together, our data suggests that ectopic NBs generated via Dpn overexpression undergo a delayed switch from neurogenesis to gliogenesis. However, they do eventually undergo terminal differentiation and exit the cell cycle, as no NBs were detected by 24 hrs APF (Figure 3I-J).

**Figure 3.**
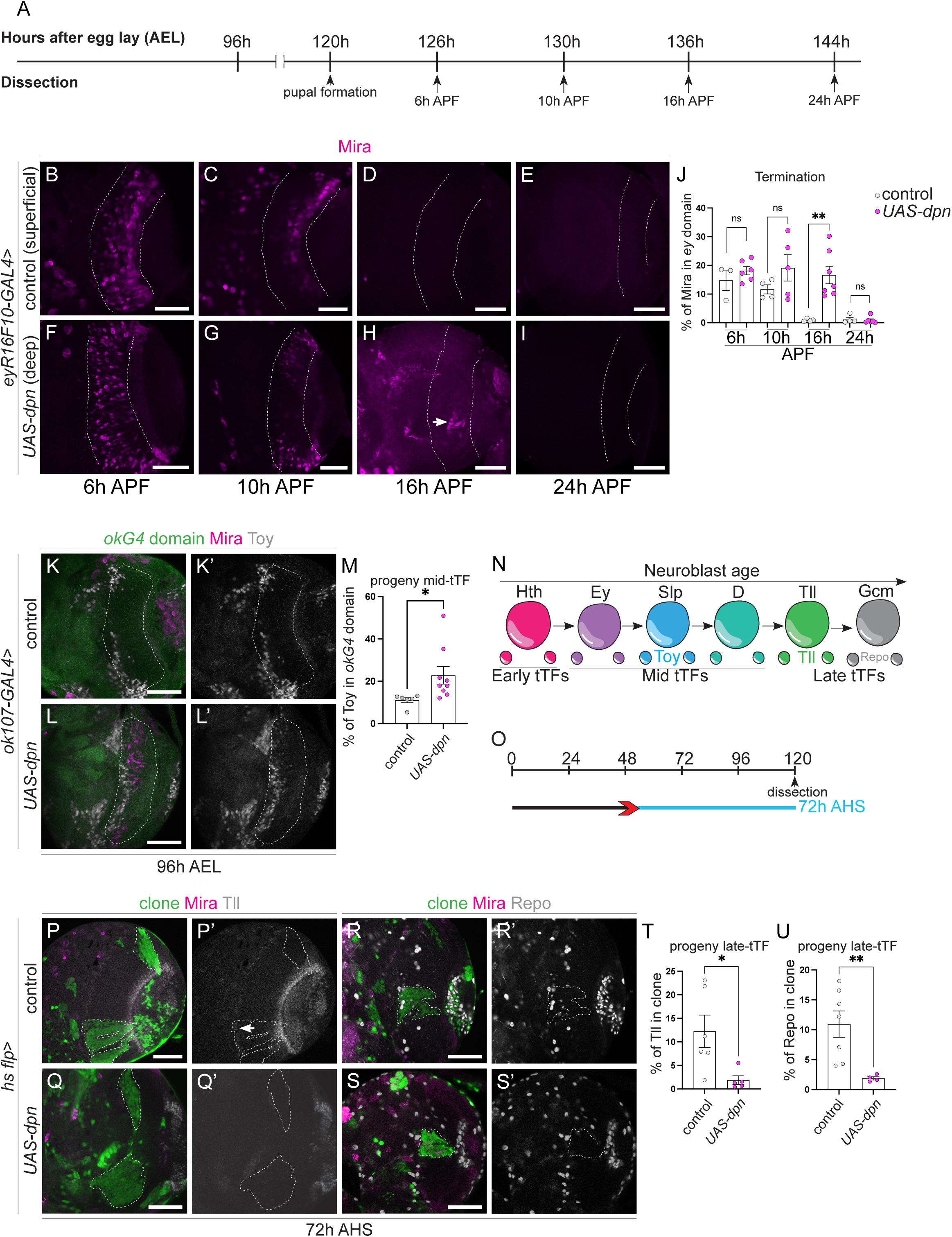
NBs generated via Dpn overexpression delays the timing of their terminal differentiation and generates Toy+ progeny at the expense of Tll+ neurons and Repo+ glial cells. (A) Schematic depicting the timeline of experiments performed in B-L’. Larval brains are dissected at 96 hr, pupal brains are dissected at 126 hr (6 hr APF), 130 hr (10 hr APF), 136 hr (16 hr APF), and 144 hr AEL (24 hr APF). (B-I, K-L’) Representative images of the superficial medulla NB layer or deep medulla neuronal layer in the larval or pupal optic lobe, in which *UAS-dpn* or control *(UAS-luciferase*) are driven by *eyR16F10-GAL4*, or *ok107-GAL4* (outlined) and stained with the stem cell marker, Miranda (Mira, magenta), and mid-temporal progeny marker, Toy (grey). (B-E) Mira+ NBs are recovered in superficial sections of control under *eyR16F10-GAL4* expression at 6 hr APF and 10 hr APF but not at 16 hr APF onwards. (F-I) Mira+ NBs are recovered in deep sections of *UAS-dpn* driven by *eyR16F10-GAL4* at 6 hr APF, 10 hr APF, and 16 hr APF but not at 24 hr APF. Arrow in G points to the NBs. (J) Quantification of % Mira+ cells within *eyR16F10-GAL4* expression domain in control and *UAS-dpn* (calculated as the ratio of Mira+ cell volume as a percentage of total *eyR16F10-GAL4* domain volume). 6 hr APF (Control, n= 3, m= 14.82 ±3.521, *UAS-dpn*, n= 6, m= 18.13 ±1.46), 10 hr APF (Control, n= 4, m= 11.65 ±1.605, *UAS-dpn*, n= 5, m= 19.11 ±4.609), 16 hr APF (Control, n= 4, m= 0.8897 ±0.239, *UAS-dpn*, n= 7, m= 16.64 ±3.065), 24 hr APF (Control, n= 4 m= 1.174, ±0.7226, *UAS-dpn*, n= 6, m= 0.8518 ±0.4849). (K-L’) More Toy+ progeny (grey) are present within the *ok107-GAL4* domain (green, outlined) in *UAS-dpn* compared to control. (M) Quantification of % Toy+ progeny within *ok107-GAL4* expression domain in control and *UAS-dpn* (calculated as the ratio of Toy+ cell volume as a percentage of total *ok107-GAL4* domain volume). Control n= 6, m= 11.03 ±1.169, *UAS-dpn*, n= 9, m= 22.78 ±4.174. (N) Schematic representation of temporal series in NBs and their progeny. NBs express temporal transcription factors (tTFs) (Hth, Ey, Slp, D, Tll) as they age. Slp+ NBs create Toy+ progeny; Tll+ NBs create Tll+ progeny; Gcm+ NBs create Repo+ progeny. (O) Heat shock regime for (P-S’). Clones were induced by heat shock (red) at 48 hr AEL and dissected 72 hr (blue) after heat shock. (P-S) Representative images of the deep medulla neuronal layer in the larval optic lobe, in which heat shock induced *UAS-dpn* or control *(UAS-luciferase*) clones are stained with the stem cell marker, Mira (magenta), late neuronal marker Tll (grey) or glial marker Repo (grey). Arrow points towards the small band of Tll+ neurons in control clones. (P-Q’) At 72 hr after clone induction, there is less Tll expression within *UAS-dpn* clones than control clone, quantified in T. (R-S’) At 72 hr after clone induction, there is less Repo expression within *UAS-dpn* clones than control clones, quantified in U. (T-U) Quantification of % Tll or Repo+ cells within control and *UAS-dpn* clones (calculated as the ratio of Tll+ or Repo+ cell volume as a percentage of total clone volume). Tll (Control n= 6, m= 12.26 ±3.449, *UAS-dpn*, n= 5, m= 1.881 ±0.9293), Repo (Control n= 7, m= 10.94 ±2.203, *UAS-dpn*, n= 4, m= 1.89 ±0.2754). Data are represented as mean ± SEM. P-values were obtained by unpaired t-test, and Welch’s correction was applied in case of unequal variances. **p<0.005, *p<0.05. Scale bars: 50 μm.

### Dpn overexpression generates Toy^+^ progeny at the expense of Tll^+^ neurons and Repo^+^ glial cells

Hth, Ey, Slp, D, and Tll are each required for the sequential generation of different medulla neurons through the production of Brain-specific homeobox (Bsh), Drifter (Dfr), Toy/Sox102F, Ets at 65A (Ets65a) and Tll, respectively (Hasegawa et al., 2011; Li et al., 2013; Morante and Desplan, 2008; Suzuki et al., 2013) (Figure 3N). We next assessed the neuronal subtypes made by Dpn-induced ectopic NBs. To better assess the cumulative effect of the neurons made throughout development, *Ey^OK107^-GAL4* was used to drive the expression of Dpn. We found almost 50% more Toy^+^ progeny was made compared to control (Figure 3 K-M). Consistent with this, *UAS-Dpn* clones produced very few Tll^+^ neurons or Repo^+^ glial cells (Figure 3 O-U). Together, our data indicates that Dpn overexpression in the medulla resulted in stalled temporal progression, producing an excess of Toy^+^ progeny at the expense of Tll^+^ neurons and Repo^+^ glial cells.

### Dichaete triggers the re-initiation of the temporal series

We have shown that Dpn overexpression induced a persistent Slp^+^ cell fate, which perturbed the generation of appropriate neuronal cell types and timely NB cell cycle exit. To investigate if Dpn activated *slp1* directly, we identified the genome-wide Dpn binding sites *in vivo* using Targeted DamID (TaDa). We profiled Dpn binding in 3rd instar neuroblasts, and identified Dpn-binding peaks at 3792 protein-coding genes. We found that Dpn directly binds to *slp1* as well as the Sox-family TF *dichaete* (*D*) which is expressed in medulla NBs after *slp1* (Li et al., 2013) (Figure S6 A-B). Consistent with the model that D represses Slp-1, we found that D and Slp were expressed in a complementary pattern in *UAS-Dpn* clones, where high expression levels of D was correlated with low expression levels of Slp, conversely low expression levels of D was correlated with high expression levels of Slp (Figure S4A-C’’’, arrows). Next, we investigated whether inhibiting Slp expression via overexpression of D was sufficient to promote temporal progression. Overexpression of D in *UAS-Dpn* clones caused a significant reduction in Slp expression, and a corresponding increase in Tll expression in NBs. Hence, D overexpression was sufficient to promote NB temporal progression towards a late temporal identity (Figure 4A-C’’, F). Consequently, D overexpression resulted in an increase in Tll^+^ neurons and Repo^+^ glial cells (Figure 4H-J’’, K, L). While a reduction in Toy^+^ progeny was also observed (Figure 4D-E’, G); this reduction was not significant, as neurons produced by D^+^ NBs can also express Toy (Bertet et al., 2014). Finally, D overexpression in *UAS-Dpn* NBs promoted their pre-mature cell cycle exit at 6 hrs APF using *eyR16F10-GAL4* (Figure 4 M-P), and flp-out clones (Figure S4 D-G).

**Figure 4.**
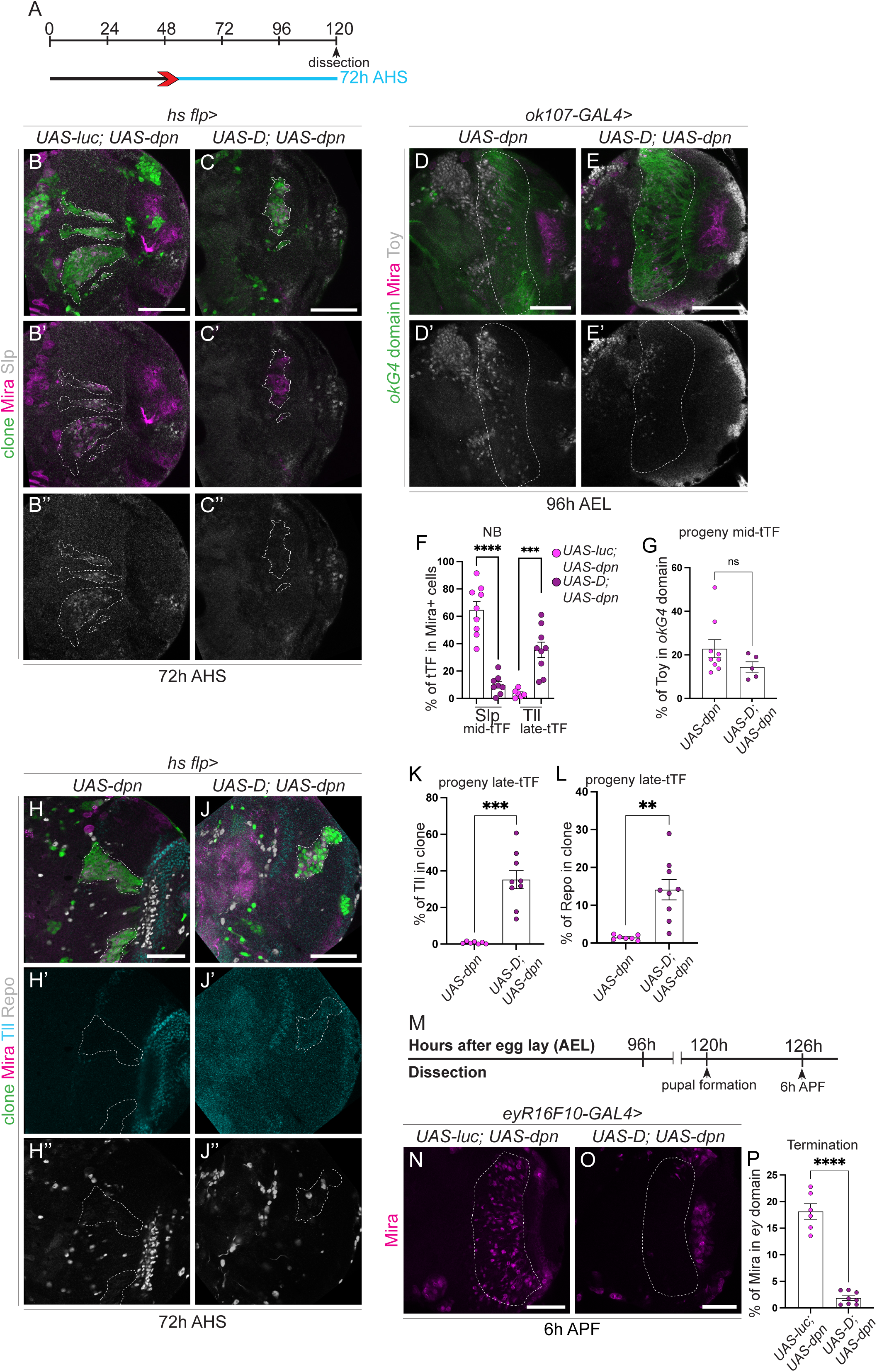
Dichaete triggers the re-initiation of the temporal series. (A) Heat shock regime for (B-C’’) and (H-J’’). Clones were induced by heat shock (red) at 48 hr AEL and dissected at 72 hr (blue) after heat shock. (B-C’’ and H-J’’) Representative images of deep medulla neuronal layer in the larval optic lobe, in which *UAS-luc*; *UAS-dpn* and *UAS-D; UAS-dpn* are expressed in clones by *hs flp*, and stained with the stem cell marker, Miranda (Mira, magenta), and/or Slp, Tll or Repo (grey). (B-C’’) At 72 hr after clone induction, Mira+ NBs within *UAS-luc; UAS-dpn* clones express Slp; Mira+ NBs within *UAS-D; UAS-dpn* clones do not express Slp, quantified in F. (D-E’) Representative images of deep medulla neuronal layer in the larval optic lobe, in which *UAS-dpn* and *UAS-D; UAS-dpn* are expressed in the *ok107-GAL4* expression domain, and stained with the stem cell marker, Mira (magenta), and mid-temporal progeny marker, Toy (grey). Fewer Toy+ cells were recovered within the *ok107-GAL4* domain (outlined) in *UAS-D; UAS-dpn* compared to *UAS-dpn*, quantified in G. (F) Quantification of % of Slp+ or Tll+ cells in Mira+ cells *UAS-luc; UAS-dpn* clones compared to *UAS-D; UAS-dpn* clones (expressed as the ratio between tTF+ volume and total Mira+ volume within a clone). Slp (*UAS-luc; UAS-dpn*, n= 9, m= 64.72 ±6.045, *UAS-D; UAS-dpn*, n= 8, m= 9.99 ±2.463), Tll (*UAS-luc; UAS-dpn*, n= 6 m= 3.584 ±1.187, *UAS-D; UAS-dpn*, n= 9, m= 35.53 ±5.596). (G) Quantification of % Toy+ cells within *ok107-GAL4* expression domain in *UAS-dpn* and *UAS-D; UAS-dpn* (calculated as the ratio of Toy+ cell volume as a percentage of total *ok107-GAL4* domain volume). *UAS-dpn* n= 9, m= 22.78 ±4.174, *UAS-D; UAS-dpn* n= 5, m= 14.42 ±2.394. (H-J’’) At 72 hr after clone induction, more Tll+ and Repo+ cells were recovered within *UAS-D; UAS-dpn* clones compared to *UAS-dpn* clones, quantified in K and L. (K-L) Quantification of % Tll+ or Repo+ cells within *UAS-dpn* clones compared to *UAS-D; UAS-dpn* clones (calculated as the ratio of Tll+ or Repo+ cell volume as a percentage of total clone volume). Tll (*UAS-dpn* n= 7, m= 0.7198 ±0.2389, *UAS-D; UAS-dpn* n= 9, m= 35.22 ±4.917), Repo (*UAS-dpn* n= 7, m= 1.488 ±0.2368, *UAS-D; UAS-dpn* n= 9, m= 14.13 ±2.672). (M) Timeline depicting the age of ectopic neuroblasts. Pupal formation occurs at 120h AEL. Pupae are dissected at 126 hr AEL (6 hr APF). (N-O) Mira+ NBs are present in deep sections of *UAS-dpn* driven by *eyR16F10-GAL4* at 126h but are absent in *UAS-D; UAS-dpn*, quantified in P. (P) Quantification of % Mira cells within *eyR16F10-GAL4* expression domain in *UAS-dpn* and *UAS-D; UAS-dpn* (calculated as the ratio of Mira+ cell volume as a percentage of total *eyR16F10-GAL4* domain volume). *UAS-dpn*, n= 6, m= 18.13 ±1.46, *UAS-D; UAS-dpn*, n= 8, m= 1.859 ±0.4215. Data are represented as mean ± SEM. P-values were obtained via unpaired t-test, and Welch’s correction was applied in cases of unequal variances. ****p<0.0001, ***p<0.001, **p<0.005. Scale bars: 50 μm.

### Temporal progression of dedifferentiated NBs is regulated by the cell cycle

Cell division is a process essential for NB development, and the progression through the cell cycle has been shown to be important for temporal transition in Type I NBs (van den Ameele and Brand, 2019). Our EdU pulse chase experiment showed that compared to control clones, *UAS-dpn* clones produced significantly fewer progeny (the number of EdU+ cells were normalised to the number of NBs), suggesting that the pace of the cell cycle was significantly slower in dedifferentiated NBs compared to that of the control (Figure 5 A-B’’, E). In addition, our DamID analysis showed that cell cycle components *cyclin D* (*cycD*), cyclin E (*cycE*) and *string* (*stg*) are direct targets of Dpn (S6 C-E). Therefore, it is possible that Dpn can directly inhibit cell cycle progression, which in turn prevents temporal transition. To test this hypothesis, we over expressed cell cycle regulators E2F transcription factor 1 (E2f1) and CycE, that are known to promote cell cycle progression at both G2/M and G1/S check points (Figure 5I). This manipulation induced a small increase in Mira^+^ NBs (Figure 5G-J) and was able to promote temporal progression, as indicated by fewer Slp^+^ NBs and a significant increase in Repo^+^ glial cells (Figure 5H-H’’’, K-L). Interestingly, overexpression of CycD and Cyclin-dependent kinase 4 (Cdk4), which promotes G1/S progression (Figure 5I), had an even stronger effect on temporal transition, resulting in premature neurogenesis to gliogenesis transition at 96 hrs AEL (Figure S5 C-E). As both D and cell cycle genes can promote temporal progression in *UAS-dpn* clones, we next assessed whether *UAS-D* also affected cell cycle progression in *UAS-dpn* clones. By performing an EdU pulse chase assay, we found that D overexpression did not significantly increase the rate of neuronal production in dedifferentiated NBs (Figure 5 C-C’’, F). Together, this data suggests that it is likely that cell cycle progression lies upstream of the temporal series, to promote the generation of neurons.

**Figure 5.**
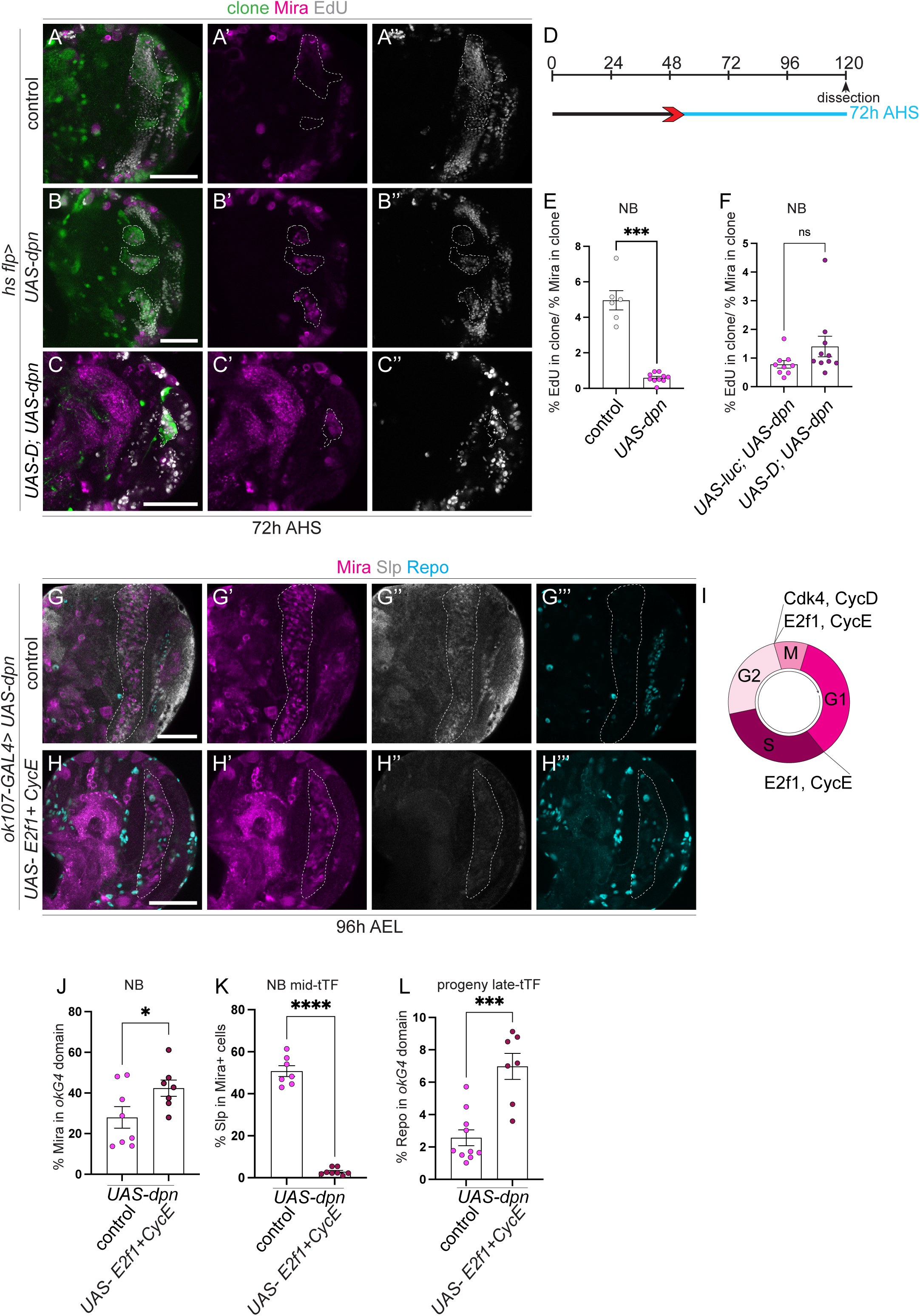
Temporal progression of dedifferentiated NBs is regulated by the cell cycle. (A-C’’) Representative images of the deep medulla neuronal layer in the larval optic lobe, in which control (*UAS-luc*) and *UAS-dpn* and *UAS-D*; *UAS-dpn* are expressed in *hs flp* clones (72 hr after clone induction). Miranda (Mira, magenta) and EdU (grey). Despite increased number of Mira+ cells in *UAS-dpn* compared to control, each NB underwent slower cell cycle progression, as indicated by EdU+ incorporation. Quantified in E. This slow cell cycle speed was not rescued in *UAS-D; UAS-dpn* clones. Quantified with *UAS-luc*; *UAS-dpn* in F. (D) Heat shock regime for (A-C’’). Clones were induced by heat shock (red) at 48 hr AEL and dissected 72 hr (blue) after heat shock. (E) Quantification of % of EdU-positive cells in control (*UAS-luc*) and *UAS-dpn* clones (expressed as the amount of EdU+ cells per Mira+ volume within the clone). Control, n= 6, m= 4.96 ±0.5433, *UAS-dpn*, n= 10, m= 0.5927 ±0.08484. (F) Quantification of % of EdU-positive cells in *UAS-luc; UAS-dpn* and *UAS-D; UAS-dpn* clones (expressed as the amount of EdU+ cells per Mira+ volume within the clone). *UAS-luc; UAS-dpn*, n= 9, m= 0.7807 ±0.1311, *UAS-D; UAS-dpn*, n= 10, m= 0.4866 ±0.3579. (G-H’’’) Representative images of the deep medulla neuronal layer in the larval optic lobe, where *UAS-E2f1; UAS-CycE*; *UAS-dpn* or *UAS-dpn* are expressed in the *ok107-GAL4* domain (outlined), and stained with the stem cell marker, Mira (magenta), mid-tTF, Slp (grey) and late progeny marker, Repo (Cyan). Quantified in J-L. *UAS-dpn* NBs express Slp and do not create Repo+ progeny, whereas *UAS-E2f1*; *UAS-CycE*; *UAS-dpn* NBs do not express Slp and create Repo+ progeny. (I) Schematic representation of the cell cycle. Cdk4/ CycD play a role in G2/M phase transition. E2f/ CycE play a role in G1/S and G2/M phase transitions. (J) Quantification of % Mira+ cells within *ok107-GAL4* expression domain in *UAS-dpn* control and *UAS-E2f1*; *UAS-CycE*; *UAS-dpn* (calculated as the ratio of Mira+ cell volume as a percentage of total *ok107-GAL4* volume). *UAS-dpn* control n= 8, m= 27.99 ±5.288, *UAS-E2f1*; *UAS-CycE*; *UAS-dpn*, n= 7, m= 42.37 ±3.999. (K) Quantification of % Slp+ cells in Mira positive cells within the *ok107-GAL4* expression domain in *UAS-dpn*; *UAS luc* and *UAS-E2f1*; *UAS-CycE*; *UAS-dpn* (calculated as the ratio of Slp+ cellular volume as a percentage of total Mira+ volume within *ok107-GAL4* domain). *UAS-dpn* control n= 7, m= 50.76 ±2.582, *UAS-E2f1*; *UAS-CycE*; *UAS-dpn*, n= 8, m= 2.907 ±0.5755. (L) Quantification of % Repo+ progeny in within the *ok107-GAL4* expression domain in *UAS-dpn*; *UAS luc* and *UAS-E2f1*; *UAS-CycE*; *UAS-dpn* (calculated as the ratio of Repo+ cellular volume within *ok107-GAL4* domain). *UAS-dpn* control n= 10, m= 2.567 ±0.491, *UAS-E2f1*; *UAS-CycE*; *UAS-dpn*, n= 7, m= 6.977 ±0.8049. Data are represented as mean ± SEM. P-values were obtained performing unpaired t-test, and Welch’s correction was applied in case of unequal variances. ****p<0.0001, ***p<0.001, **p<0.005, *p<0.05. Scale bars: 50 μm.

### Notch-mediated dedifferentiation also causes temporal series progression defects

To look at whether dedifferentiated NBs commonly exhibit defects in their temporal identity, we next examined another model of neuronal dedifferentiation. We have previously identified Notch (N) to be a Nerfin-1 target gene, where N activation phenocopied loss of Nerfin-1, resulting in neuron to NB dedifferentiation (Vissers et al., 2018) (Figure 6 A-E). Recently, it was shown that the N pathway regulates Slp1 and Slp2 expression in medulla NBs through direct binding to Slp1/2 enhancers (Ray and Li, 2021). To test whether similar to Dpn overexpression, N activation also stalled tTF progression, we assessed Slp expression in medulla regions where *UAS-N^ACT^* was expressed under the control of *eyR16F10-GAL4*. We found that the ectopic NBs induced via N^ACT^ exhibited significantly increased expression of mid-tTF Slp (Figure 6C-C’’’, F), consequently resulting in an increase in the number of Toy^+^ neurons (Figure 6I-K). Similar to Dpn overexpression, dedifferentiated NBs caused by N activation failed to undergo timely terminal cell cycle exit. At 24 hrs APF, where all of the wildtype NBs have exited the cell cycle, NBs generated via N activation continued to persist (Figure S5A-B).

**Figure 6.**
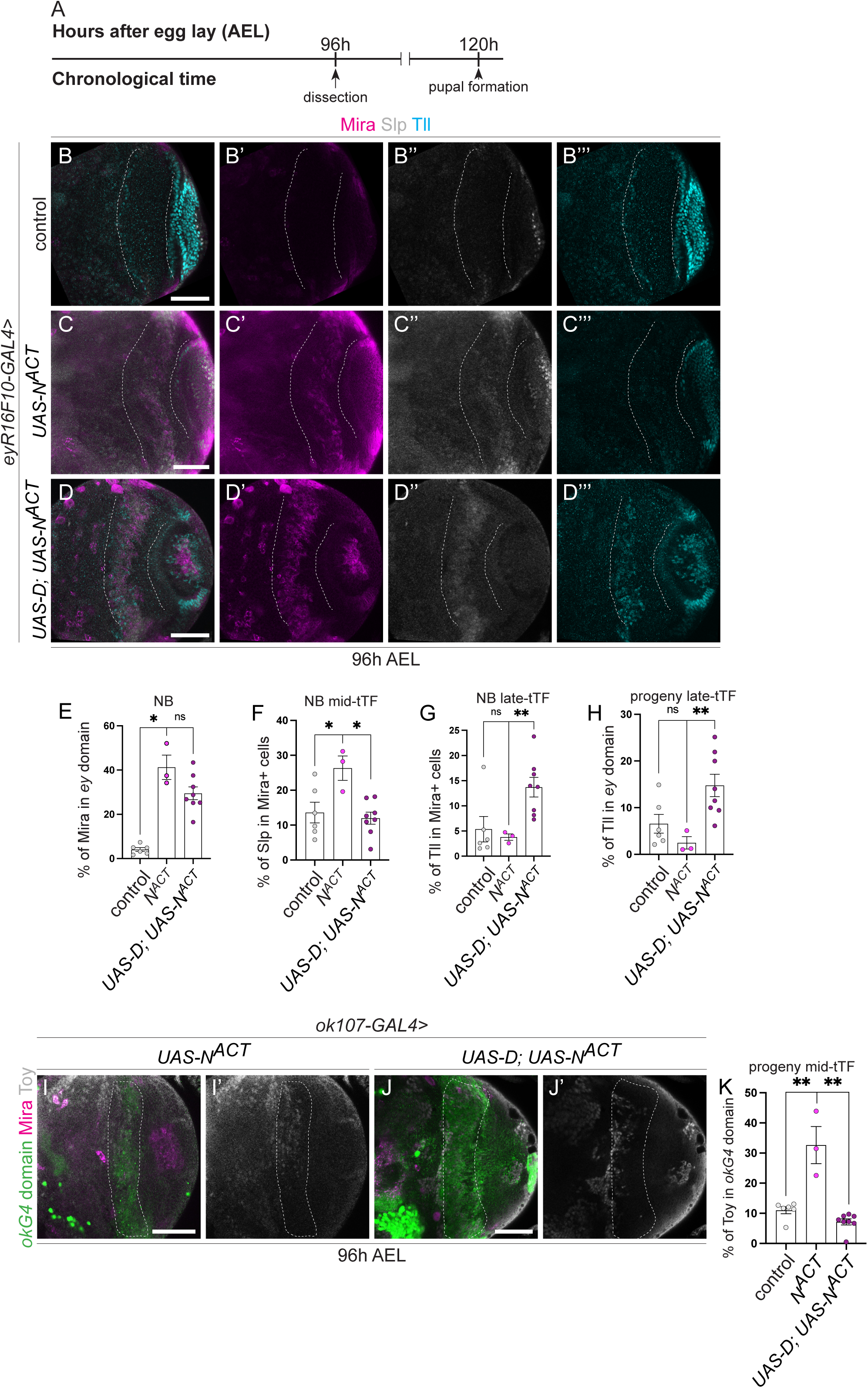
Dichaete re-initiates the temporal series in dedifferentiated NBs caused by Notch activation. (A) Timeline depicting age of ectopic neuroblasts. Larvae are dissected at 96h AEL. (B-D’’’) Representative images of the deep medulla neuronal layer in the larval optic lobe, in which control (*UAS-luc*), *UAS-N^ACT^*, and *UAS-D; UAS-N^ACT^* are driven by *eyR16F10-GAL4* (outlined). Miranda (Mira, magenta), Slp (grey), Tll (blue). *UAS-N^ACT^* NBs express Slp and not Tll. *UAS-D; UAS-N^ACT^* NBs express Slp and Tll, quantified in E-H. (E) Quantification of % Mira-positive cells within *eyR16F10-GAL4* expression domain in control, *UAS-N^ACT^*, and *UAS-D; UAS-N^ACT^* (calculated as the ratio of Mira+ cell volume as a percentage of total volume of *eyR16F10-GAL4* domain). Control n= 6, m= 4.077, ±2.103, *UAS-N^ACT^* n= 3, m= 41.26, ±5.502, *UAS-D*; *UAS-N^ACT^* n= 8, m= 29.5, ±2.931. (F-G) Quantification of % Slp+ or Tll+ cells in Mira positive cells within the *eyR16F10-GAL4* expression domain in control, *UAS-N^ACT^*, and *UAS-D; UAS-N^ACT^* (calculated as % of Slp+ or Tll+ cellular volume that are also Mira+ within *eyR16F10-GAL4* domain). Slp (Control n= 6, m= 13.62 ±2.974, *UAS-N^ACT^* n= 3, m= 26.34 ±3.47, *UAS-D*; *UAS-N^ACT^* n= 8, m= 11.98 ±1.741), Tll (Control n= 6, m= 5.363 ±2.531, *UAS-N^ACT^* n= 3, m= 3.775 ±0.6369, *UAS-D*; *UAS-N^ACT^* n= 8, m= 13.71 ±1.955). (H) Quantification of % Tll+ cells within *eyR16F10-GAL4* expression domain in control, *UAS-N^ACT^*, and *UAS-D; UAS-N^ACT^* (calculated as the ratio of Tll+ cellular volume as a percentage of total volume of *eyR16F10-GAL4* domain). Control n= 6, m= 6.554 ±2.018, *UAS-N^ACT^* n= 3, m= 2.457 ±1.323, *UAS-D*; *UAS-N^ACT^* n= 8, m= 14.77 ±2.404. (I-J’) Representative images of the deep medulla neuronal layer in the larval optic lobe, in which *UAS-N^ACT^*and *UAS-N^ACT^; UAS-dpn* are driven by *ok107-GAL4* (marked by GFP and outlined) and stained with the stem cell marker, Mira (magenta), and mid-temporal progeny marker, Toy (grey). Fewer cells express Toy+ in *UAS-N^ACT^; UAS-dpn* compared to *UAS-N^ACT^*, quantified in K. (J) Quantification of % Toy+ cells within *ok107-GAL4* expression domain in control (*UAS-luc*), *UAS-N^ACT^* and *UAS-N^ACT^; UAS-dpn* (calculated as the ratio of Toy+ cell volume as a percentage of total *ok107-GAL4* domain volume). The column for Control uses the same data points as in Figure 3K. Control n= 6, m= 11.03 ±1.169, *UAS-N^ACT^*, n= 3, m= 32.65 ±6.176, *UAS-N^ACT^; UAS-dpn*, n= 8, m= 7.178 ±1.023). Data are represented as mean ± SEM. P-values were obtained performing unpaired t-test, and Welch’s correction was applied in case of unequal variances. ****p<0.0001, ***p<0.001, **p<0.005. Scale bars: 50 μm.

Next, we asked whether overexpression of D, known to repress Slp, could re-initiate temporal progression, restore neuronal progeny composition and timely terminal differentiation in Notch mediated dedifferentiation. Overexpression of D did not significantly affect the rate of dedifferentiation (Figure 6E), but significantly increased the proportion of NBs that expressed the late tTF Tll, and reduced the proportion of NBs that expressed the mid tTF Slp (Figure 6 F-G). Consequently, this also resulted in an increase in the amount of Tll^+^ neurons and a reduction in Toy^+^ progeny (Figure 6D-D’’’, H-K). Together, our data suggest that D overexpression can promote temporal progression stalled via Dpn overexpression or Notch activation.

## Discussion

In this study, our data suggests that upon dedifferentiation, the proliferative potential and ability to produce the correct neuronal subtypes is determined by cell cycle progression and the temporal transcription factors. We showed that upon Dpn overexpression, ectopic NBs were induced. These ectopic NBs immediately adopted a Slp^+^ mid-temporal identity. Consequently, this significantly increased the amount of mid-temporal window progeny produced compared to control, slowed down the cell cycle and delayed terminal cell cycle exit. Re-triggering the temporal series via D overexpression as well as promoting the cell cycle were sufficient to reinstate neuronal diversity and timely termination (Figure 7). This mechanism also held true for ectopic NBs induced via N hyperactivation.

**Figure 7.**
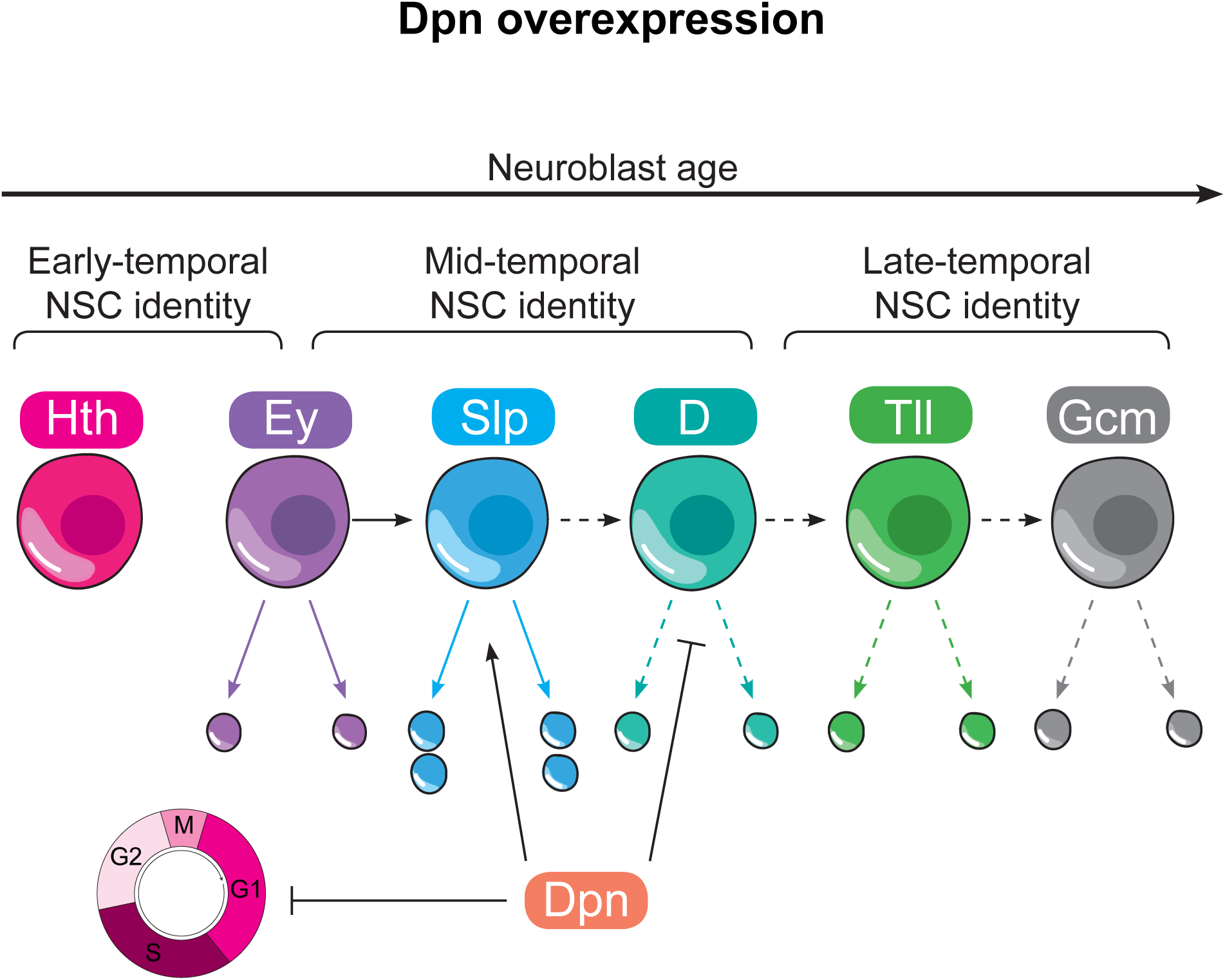
Dpn overexpression leads to delayed temporal progression via cell cycle regulation. Upon Dpn overexpression, ectopic NBs were induced. These ectopic NBs expressed the mid-temporal tTF, Slp, and did not express early tTFs or late tTFs. Consequently, an over-abundance of progeny was produced from the Slp^+^ temporal window at the expense of the progeny produced from the late time windows. Slp, D, and cell cycle genes are targets of Dpn; therefore, Dpn may directly be responsible for the increase in Slp, decrease in D and stalled cell cycle progression in dedifferentiated NBs.

Our data supports the possible links between cell cycle progression and the expression of temporal regulators controlling NB proliferation and cellular diversity. In the embryonic NBs, temporal transition can progress despite of cell cycle arrest (Grosskortenhaus et al., 2005). Whereas in Type I larval NBs, it was shown that prolonged G1/S-phase resulted in delayed temporal patterning and reduced neuronal diversity (van den Ameele and Brand, 2019). In dedifferentiated NBs, it appears that the temporal series lies downstream of the cell cycle. Overexpression of cell cycle regulators can promote temporal progression and restore neuronal diversity, but re-initiation of the temporal series was not sufficient to rescue cycle cycle progression. Cell cycle therefore directs the progression of the temporal series in this particular context.

In addition to cell cycle regulation, we found that temporal transitions in dedifferentiated NBs also obey feed forward activation and feedback repression that occurs in wildtype medulla NBs. We found D is expressed in a complementary pattern to that of Slp-1, and D overexpression can re-trigger the progression of the temporal series (Li et al., 2013). Interestingly, cell cycle and temporal transcription factors were all direct target genes of Dpn. It would be interesting to explore whether N similarly acts on these target genes to specify cell fate and proliferation profiles of dedifferentiated NBs. However, it could be that N acts partially through Dpn, as Dpn has been shown to be a target of N in NBs (San-Juán et al., 2012).

Recent studies suggest that a bidirectional conversion between tumour-initiating stem cells and differentiated cells exist, where non-cancer stem cells can dedifferentiate and acquire stem cell-like properties under genetic and environmental stress (Friedmann-Morvinski and Verma, 2014). Dedifferentiation of mature cells to induce their proliferation and redifferentiation, has also been shown to be a promising means of tissue repair and regeneration. Therefore, dedifferentiation may have either positive or negative consequences for tissue repair and tumorigenesis. Our work shows that appropriate temporal and cell cycle progression controls are key to regulating the fine balance between tumorigenesis and controlled dedifferentiation.

## Methods and materials

### Fly husbandry

Fly stocks were reared on standard *Drosophila* media at 25 °C. For larval dissection, brains were dissected at wandering L3 stages. For pupal dissection, white pupae were selected and then allowed to age at 29 °C, dissections were made at 6, 10, 16, or 24 hours after pupal formation (APF). Pupal experiments were generated using *eyR16F10-GAL4* unless otherwise stated. Additional drivers used for larval experiments were: *GMR31H08-Gal4* (Jenett et al., 2012)*, UAS-GFP/ TM6b*; *ey^OK107^-GAL4* (Morante et al., 2011).

Knock-down or overexpression CNS clones were generated using the FLIPOUT system. *hsFLP; actc5>CD2>Gal4, UAS-GFP/ TM6b* (Vissers et al., 2018). FLIPOUT clones were induced by heat shock at 16, 24, 48h, 72, or 96 hr after egg lay (AEL) for 6 min, shifted to 29°C, and dissected at 120h AEL. The fly strains used were: *UAS-deadpan* (A. Baonza), (#1776, Bloomington), *UAS-Nerfin-1 RNAi* (VDRC), *UAS-mCherry RNAi* (#35785, Bloomington), *Toy-GFP* (#83390, Bloomington), *UAS-N^ACT^*(#52309, Bloomington), *UAS-Cdk4; UAS-CycD* (Helena Richardson), *UAS-E2f1; UAS-CycE* (Helena Richardson), *UAS-D* (#8861, Bloomington),.

### Immunostaining

Larval and pupal brains were dissected out in phosphate buffered saline (PBS), fixed for 20 minutes in 4% formaldehyde in PBS and rinsed in 0.2% PBST (PBS + 0.2% TritonX-100). For immunostaining, brains were incubated with primary antibodies overnight at 4°C, followed by an overnight secondary antibody incubation at 4°C. Samples were mounted in 80% glycerol in PBS for image acquisition. The primary antibodies used were mouse anti-Mira (1:50; gift of Alex Gould), rat anti-Mira (1:100, Abcam), rat anti-pH3 (1:500; Abcam), chick anti-GFP (1:2000; Abcam), rabbit anti-RFP (1:100, Abcam), mouse anti-Repo (1:50, DSHB), guineapig anti-Hth (1:100, Claude Desplan), mouse anti-Ey (1:50, DSHB), guineapig anti-Slp (1:500, Kuniaki Saito), rabbit anti-D (1:1000, Steve Russell), guineapig anti-Tll (1:100, Kuniaki Saito), rabbit anti-Tll (1:100, Kuniaki Saito), and guineapig anti-Toy (1:50, Uwe Walldorf). Secondary donkey antibodies conjugated to Alexa 555 and Alexa 647, and goat antibody conjugated to Alexa 405, 488, 555 and 647 (Molecular Probes) were used at 1:500.

### EdU pulse/chase

Control and *UAS-d*pn clones were induced 48 hrs AEL. 48 hrs after clone induction, larvae were fed with 100 mg/mL EdU for 3 hrs. They were then transferred to standard medium for a 3 hrs EdU-free chase. For experiments using *GMR31H08-Gal4* driver (Figure S2): 48h AEL larvae were fed with 100mg/mL EdU for 2 hours and transferred to a standard medium for a 2 hrs EdU-free chase. Larvae were dissected, fixed and processed for antibody staining, followed by EdU detection with Click-iT Plus EdU Cell Proliferation Kit for imaging, Alexa Fluor 647 dye (Invitrogen, #C10640) according to the manufacturer’s instruction.

### Imaging and image processing

Images were collected on a Leica SP5/ Olympus FV3000 confocal microscope and processed using Fiji (https://imagej.net/Fiji). Z stacks of optic lobes were imaged and single sections were shown as the representative image in the figures.

### Statistical analysis

At least three animals per genotype were used for all experiments. Volume of clones or regions of interest was estimated from three-dimensional reconstructions of 2-mm spaced confocal Z stacks with Volocity software (Improvision).

Clone volume was calculated by making a Region Of Interest (ROI) around the GFP^+^ clone (excluding the top section where the superficial NBs are). *GMR31H08-Gal4* and *ey^OK107^-GAL4* expressing domains were labelled with RFP or GFP, and the overall volume of the ROI was calculated as above. *eyR16F10-GAL4* domain was not labelled with a fluorescent marker, therefore a ROI was manually selected as guided by the outline of the overlaying superficial NBs.

The rate of dedifferentiation was represented as the volume of Mira^+^ cells as a percentage of clone volume or domain volume. To estimate the percentage of control or dedifferentiated NBs that expressed a specific tTF, we made a ROI from Mira^+^ cells within the clone or the domain of interest, and selected for voxels that were positive for the specific tTF within this ROI.

To estimate the percentage of progeny that expressed a specific tTF, we made a ROI from the clone or the expression domain, and selected for cells that were positive for the specific tTF within this ROI. In all graphs and histograms, error bars represent the standard error of the mean (SEM) and p values are calculated by performing two-tailed, unpaired Student’s t test. The Welch’s correction was applied in case of unequal variances.

### Targeted DamID

FlyORF-TaDa-Dpn flies were created from the FlyORF-Dpn-3xHA line (FlyORF stock# F000086) with the FlyORF-TaDa system as previously described (Aughey et al., 2021). Targeted DamID for Dpn on 3rd instar larval neuroblasts was performed as previously described (Marshall and Brand, 2017; Marshall et al., 2016), crossing FlyORF-TaDa-Dpn flies with the neuroblast-specific w;wor-Gal4;tub-GAL80ts driver. Briefly, flies were allowed to lay on apple juice plates with yeast for 4 hours at 25°C, before transferring plates to 18°C for two days. 100 larvae from each plate were transferred to food plates and grown at 18°C for a further five days, before shifting to the permissive temperature of 29°C. Two biological replicates were collected, of thirty dissected brains per replicate.

Samples were processed for DamID as previously described (Marshall et al., 2016). Briefly, genomic DNA was extracted, cut with DpnI and cut fragments isolated. DamID adaptors were ligated to the isolated DNA, fragments were digested with DpnII, and amplified via PCR. Following PCR, samples were sonicated in a Bioruptor Plus (Diagenode) to reduce the average DNA fragment size to 300bp, and DamID adaptors were removed via overnight AlwI digestion. The resulting DNA was purified via magnetic bead cleanup and 500ng of DNA was end-repaired, A-tailed, ligated to NGS adaptors and amplified via 6 PCR cycles. The final libraries, multiplexed to yield ~20 million mappable fragments per sample, were sequenced as paired-end reads (MGI platform, BGI).

NGS reads in FASTQ format were aligned using damidseq_pipeline (Marshall and Brand, 2015) using default options, and replicates were averaged to give the final binding profile. Significantly-enriched peaks were called on this final binding track using find_peaks (Marshall et al., 2016) with parameters --min-quant=0.8 and --unified_peaks=min with otherwise default options; peaks were considered significant at FDR<0.01.

Datasets were visualised using pyGenomeTracks (Cubeñas-Potts et al., 2017).

## Acknowledgements

We are grateful to C. Desplan, H. Richardson, S.Russell, U.Walldorf, K Saito, for generous sharing of antibodies and fly stocks. L.Y.C is funded by an Australian Research Council Future Fellowship, L.Y.C’s laboratory is supported by funding from the NHMRC, ARC and the Peter MacCallum Cancer Foundation. Confocal Microscopy was performed at the Centre for Advanced Histology & Microscopy at the Peter MacCallum Cancer Centre and the Biological Optical Microscopy Platform at the University of Melbourne.

## Author contribution

K.V., F.F., Q.D., E.A., K.N, O.M, J.P.D.M and L.Y.C. conducted the experiments; K.H and P.R.J provided intellectual input, K.V and L.Y.C. designed the experiments and wrote the paper.

---

**Supplementary 1.**
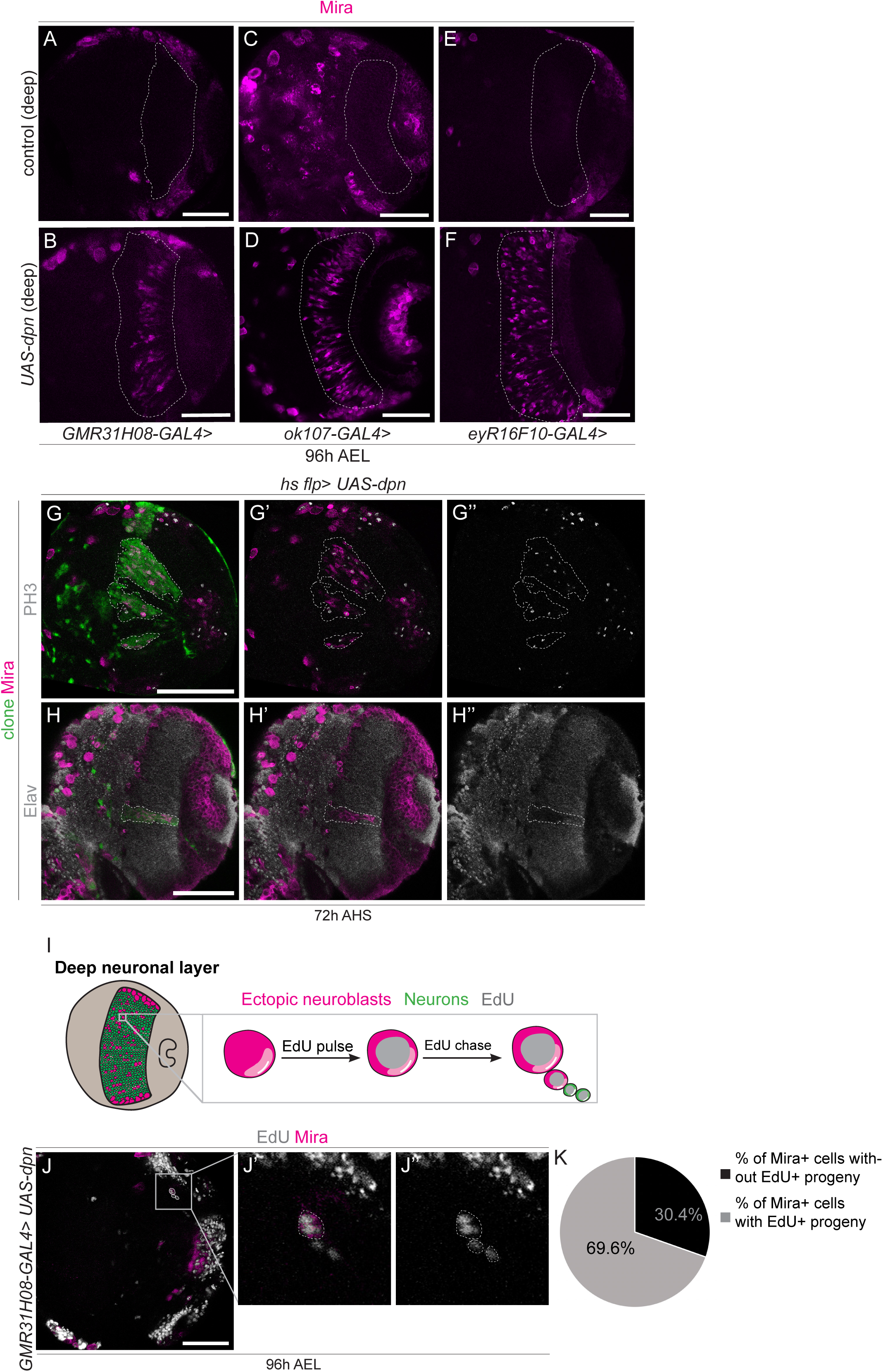
Dpn overexpression causes dedifferentiation in deep layers of the OL. (A-F) Representative images of the deep medulla neuronal layer in the larval optic lobe, in which *UAS-dpn* and control are expressed using *GMR31H08-GAL4, eyR16F10-GAL4*, and *ok107-GAL4* (outlined) and stained with the stem cell marker, Miranda (Mira, magenta). (G-H’’) Representative images of the deep medulla neuronal layer in the larval optic lobe, in which *UAS-dpn* and control clones are generated via *hs flp*. At 72h after *UAS-dpn* clone induction, Mira+ NBs (magenta) express M phase marker PH3 (grey) and do not express neuronal marker Elav (grey). (I) Schematic representation of ectopic NBs present in deep neuronal layers of the medulla. Larvae are fed EdU-labelled food for 2 hours and chased for 2 hours with an EdU-free food. Neuronal progeny (green) generated by ectopic dedifferentiated NBs (pink) would inherit EdU (grey), but would not be Mira+. (J’-J’’) are magnified images of (J). Two Mira-negative, EdU+ progeny are produced by a ectopic NBs. Here, *UAS-dpn* expression is driven by *GMR31H08-GAL4*, quantified in K. (K) Quantification of % of Mira+ ectopic NBs with or without EdU+ progeny. n=46 Scale bars: 50 μm.

**Supplementary 2.**
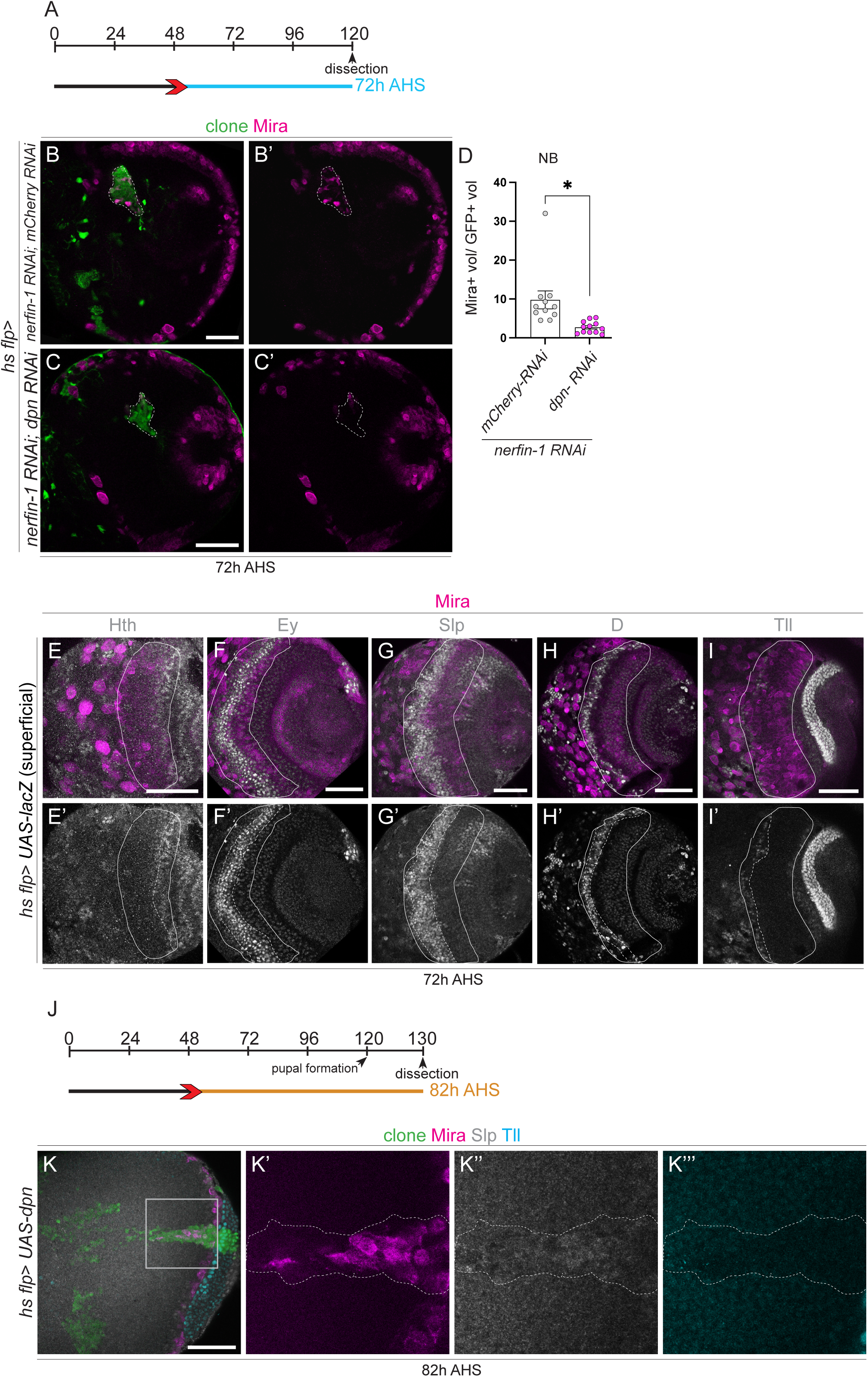
Dpn is epistastic to Nerfin-1 RNAi, representative examples of superficial section of the OL expressing tTFs and UAS-Dpn clones express Slp during pupae neurogenesis. (A) Heat shock regime for (B-C). Clones were induced by heat shock (red) at 48 hr AEL and dissected 72 hr (blue) after heat shock. (B-C) Representative images of the deep medulla neuronal layer in the larval optic lobe, in which *UAS-dpn*, *nerfin-1 RNAi*; *mCherry RNAi*, *nerfin-1 RNAi; dpn RNAi* are driven in clones by *hs flp* (marked by GFP and outlined) and stained with the stem cell marker, Miranda (Mira, magenta), quantified in D. (D) Quantification of % Mira+ cells in control *nerfin-1 RNAi; mCherry RNAi* and *nerfin-1 RNAi; dpn RNAi* clones (calculated as the ratio of Mira+ cell volume as a percentage of total clone volume). *nerfin-1 RNAi; mCherry RNAi* n=11, m=9.774 ±2.325, *nerfin-1 RNAi; dpn RNAi* n=12, m=2.722 ±0.441. (E-I’) Representative images of *hs flp* induced clones, where the Mira+ NBs within the clone in the superficial section of the medulla expressing the temporal transcription factors (Hth, Ey, Slp, D, Tll). (I) Heat shock regime for (K-K’’’). Clones were induced by heat shock (red) at 48 hr AEL and dissected at 126 hr AEL (6 hr APF), corresponding to 82 hr (blue) hours after heat shock. (K-K’’’) are magnified images of (K). At 82 hr after clone induction, Mira+ NBs express Slp and not Tll. Data are represented as mean ± SEM. P-values were obtained performing unpaired t-test, and Welch’s correction was applied in case of unequal variances. *p < 0.05. Scale bars: 50 μm.

**Supplementary 3.**
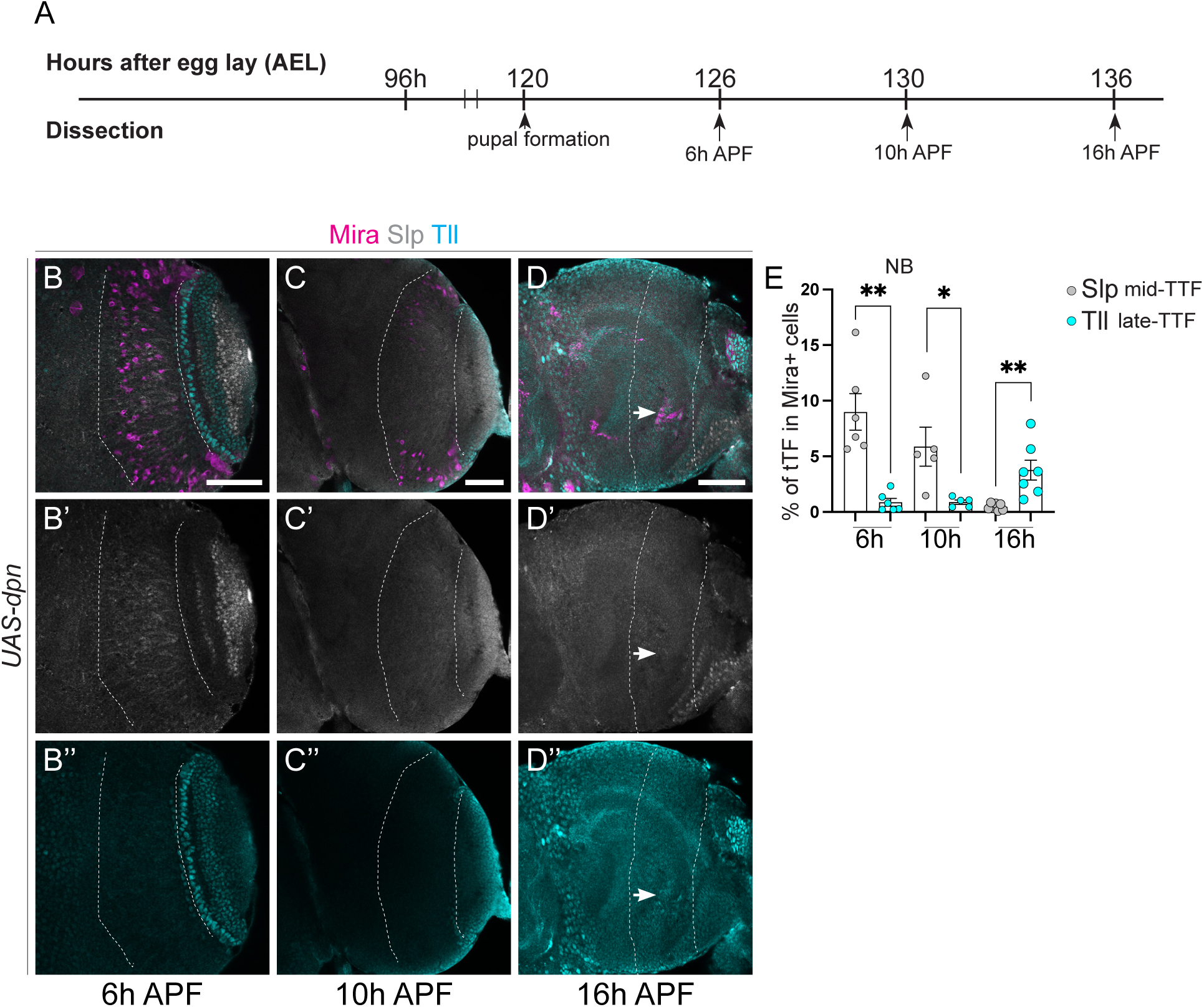
Slp and Tll expression in dedifferentiated NBs induced via Dpn overexpression during pupal stages. (A) Time-line depicting age of ectopic neuroblasts. Pupal formation occurs at 120h AEL; pupae are dissected at 126 hr (6 hr APF), 130 hr (10 hr APF), AND 136 hr (16 hr APF). (B-D’’) Representative images of the deep medulla neuronal layer in the pupal optic lobe, in which *UAS-dpn* is driven by *eyR16F10-GAL4* (outlined) and stained with the stem cell marker, Miranda (Mira, magenta). Mid-tTF Slp and late neuronal marker Tll (cyan). Dedifferentiated NBs express Slp and do not express Tll at 6 hr APF and 10 hr APF; however, they by 16 hr APF they do not express Slp and instead express Tll. Quantified in E. (E) Quantification of % Slp+ or Tll+ cells in Mira positive cells within the *eyR16F10-GAL4* expression domain in *UAS-dpn* (calculated as the ratio of Slp/Tll+ cell volume and Mira+ volume within *eyR16F10-GAL4* domain). 6 hr APF (Slp n= 6, m= 8.999 ±1.64, Tll n= 6, m= 0.8718, ±0.3451). 10 hr APF (Slp n= 5, m= 5.876, ±1.752, Tll n= 5, m= 0.89 ±0.1915). 16 hr APF (Slp n= 7, m= 0.4699, ±0.116, Tll n= 7, m= 3.767 ±0.8893). Data are represented as mean ± SEM. P-values were obtained performing unpaired t-test, and Welch’s correction was applied in case of unequal variances. **p<0.005, *p<0.05. Scale bars: 50 μm.

**Supplementary 4.**
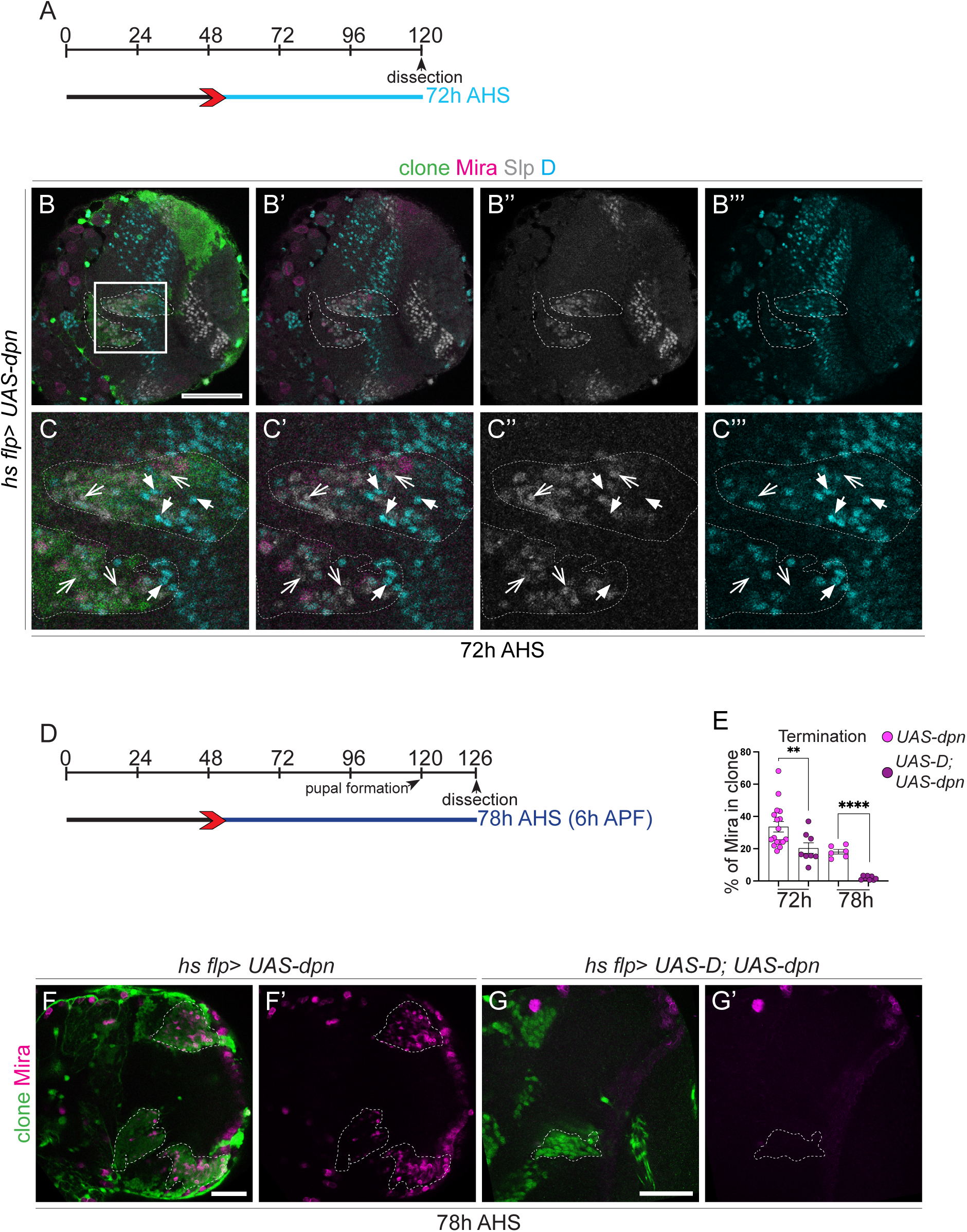
Slp and D are expressed in complementary patterns and UAS-D promotes premature termination of *UAS-dpn* ectopic NBs. (A) Schematic depicting the heat shock regimes used in (B-C’’). Clones were induced via heat shock (red arrows) at 48 hr and dissected 72 hr (blue) after heat shock. (B-C’’) Representative images of the deep medulla neuronal layer in the larval optic lobe, where *UAS-dpn* generated by *hs flp*, express complementary patterns of Slp and D expression. (C-C’’’) are magnified of (B-B’’’). At 72 hr after clone induction, Mira+ NBs express Slp or D. Open arrow heads represent NBs that express Slp and not D. Closed arrow heads represent NBs that express D and not Slp. Miranda (Mira, magenta), D (cyan), Slp (grey). (D) Schematic depicting the heat shock regimes used in (F-G’). Clones were induced via heat shock (red arrows) at 48 hr and dissected at 78 hr (dark blue) after heat shock. Pupal formation occurs at 120 hr AEL; 78 hr after heat shock is equivalent to is 6 hr APF. (E) Quantification of % Mira+ cells in control *UAS-dpn* and *UAS-D; UAS-dpn* clones (calculated as the ratio of Mira+ cell volume as a percentage of total clone volume). 72 hr (*UAS-dpn* n=17, m=33.62 ±3.243, *UAS-D; UAS-dpn* n=8, m=20.38 ±3.307). 78 hr (*UAS-dpn* n=6, m=18.3 ±1.46, *UAS-D; UAS-dpn* n=8, m=1.859 ±0.4215). (F-G’) Representative images of the deep medulla neuronal layer in the larval optic lobe, where *UAS-dpn* or *UAS-D; UAS-dpn* clones are inducted via *hs flp*. NBs are marked by stem cell marker, Mira (magenta). At 72 hr and 78 hr (6 hr APF) after clone induction, NBs underwent premature terminal differentiation upon overexpression of D, quantified in E. Data are represented as mean ± SEM. P-values were obtained performing unpaired t-test, and Welch correction was applied in case of unequal variances. ****p<0.0001, **p<0.005. Scale bars: 50 μm.

**Supplementary 5.**
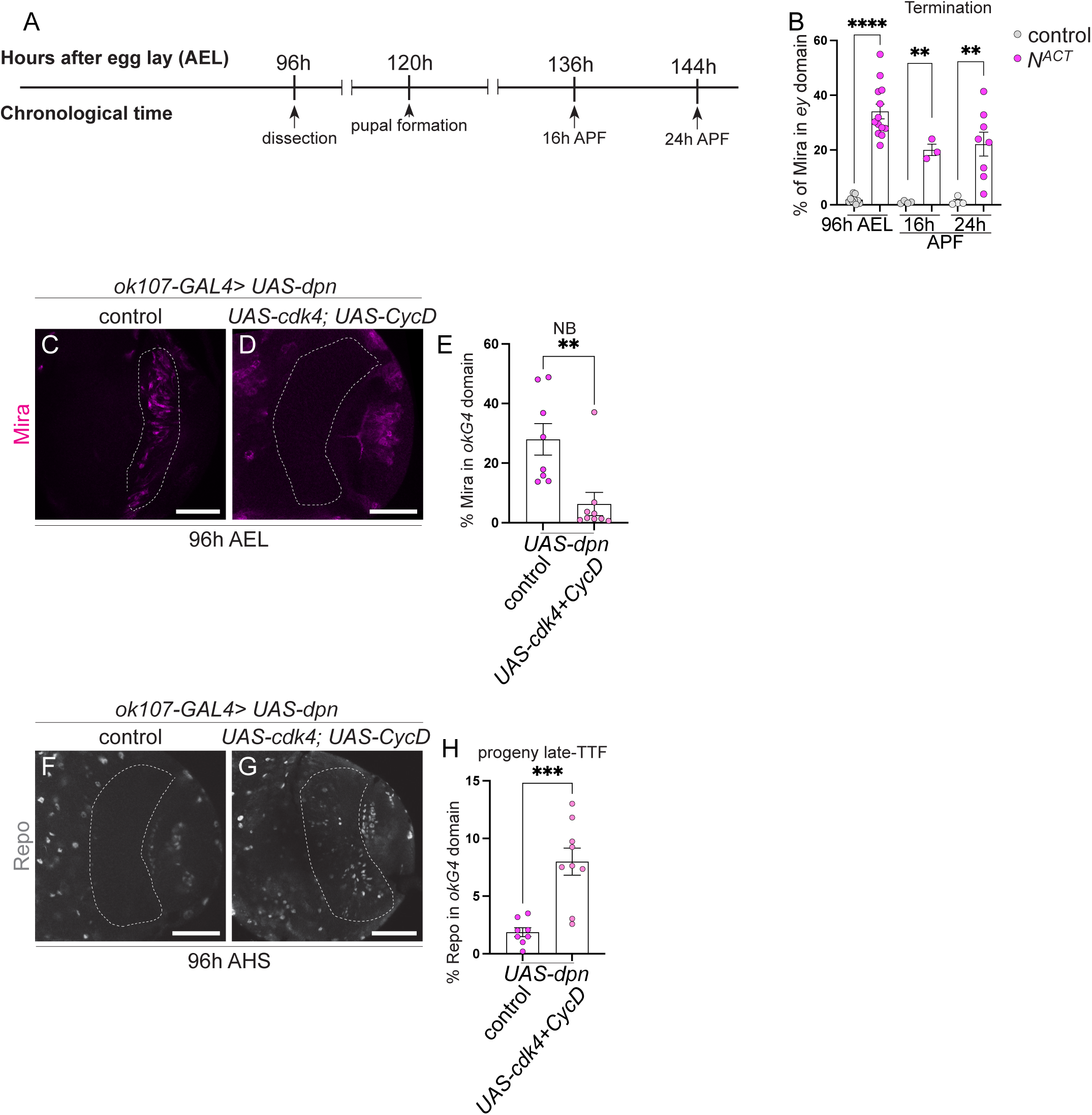
N^ACT^ ectopic NBs do not terminate at 24 hr APF and Cdk4+CycD promote temporal progression in dedifferentiated NBs induced via Dpn overexpression. (A) Time line depicting age of ectopic neuroblasts. Larvae are dissected at 96 hr. Pupal formation occurs at 120 hr AEL; pupae are dissected at 136 hr (16 hr APF) and 144 hr AEL (24 hr APF). (B) Quantification of % Mira+ cells within *ok107-GAL4* expression domain in *UAS-N^ACT^* and control (calculated as the ratio of Mira+ cell volume as a percentage of total *ok107-GAL4* domain volume). 96 hr (control n= 9, m= 1.874 ±0.4897, *UAS-N^ACT^* n= 13, m= 34.06, ±2.7). 136 hr (control n= 4, m= 0.8897, ±0.239, *UAS-N^ACT^* n= 3, m= 20.04 ±2.091). 144 hr (control n= 4, m= 1.174 ±0.7226, *UAS-N^ACT^* n= 8 m= 22.14, ±4.382). (C-D) Representative images of the deep medulla neuronal layer in the larval optic lobe, where *UAS-cdk4; UAS-CycD*; *UAS-dpn* or *UAS-dpn; UAS luc* are expressed in the *ok107-GAL4* domain (outlined), and stained with the stem cell marker, Mira (magenta). (E) Quantification of % Mira-positive cells within *ok107-GAL4* expression domain in *UAS-dpn* control and *UAS-cdk4; UAS-CycD*; *UAS-dpn* (calculated as the ratio of Mira+ cell volume as a percentage of total *ok107-GAL4* domain volume). *UAS-dpn* control n= 8, m= 27.99, ±5.288, *UAS-cdk4; UAS-CycD*; *UAS-dpn*, n= 9, m= 6.275, ±3.906. (F-G) Representative images of the deep medulla neuronal layer in the larval optic lobe, where *UAS-dpn* control or *UAS-cdk4; UAS-CycD; UAS-dpn* are expressed in the *ok107-GAL4* domain. Glial cells are labelled with Repo (grey) within the *ok107-GAL4* domain (outlined). Quantified in H. (H) Quantification of % Repo-positive cells within *ok107-GAL4* expression domain in *UAS-dpn* and *UAS-cdk4; UAS-CycD; UAS-dpn* (calculated as the ratio of Repo+ cell volume as a percentage of total *ok107-GAL4* domain volume). *UAS-dpn* n= 8, m= 1.875, ±0.3837, *UAS-cdk4; UAS-CycD; UAS-dpn*, n= 9, m= 7.997, ±1.173. Data are represented as mean ± SEM. P-values were obtained performing unpaired t-test, and Welch correction was applied in case of unequal variances. ****p<0.0001, ***p<0.001, **p<0.005, *p<0.05. Scale bars: 50 μm.

**Supplementary 6.**
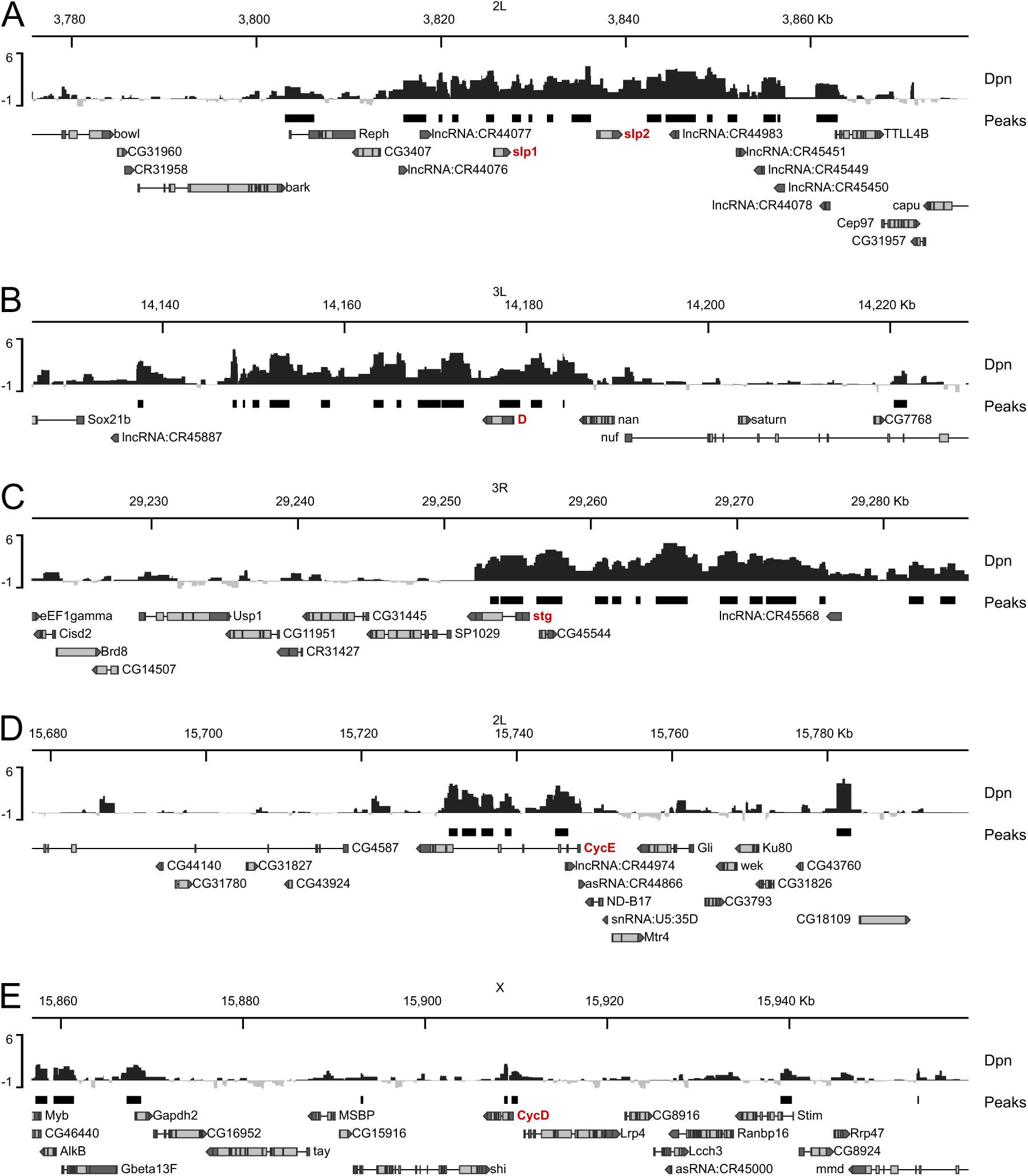
Dpn binds to tTFs and cell cycle loci. Mean Targeted DamID Dpn binding profiles (log2(Dam-fusion/Dam)) are shown for the optic lobe temporal TFs (A) slp1 / slp2 and (B) D; and for the cell cycle genes (C) stg, (D) CycE and (E) CycD. Significantly enriched binding peaks overlapped with each of the five gene loci; peaks with FDR<0.01 are shown.

